# Protein-lipid interaction at low pH induces oligomerisation of the MakA cytotoxin from *Vibrio cholerae*

**DOI:** 10.1101/2021.09.10.459761

**Authors:** Aftab Nadeem, Alexandra Berg, Hudson Pace, Athar Alam, Eric Toh, Jörgen Ådén, Nikola Zlatkov, Si Lhyam Myint, Karina Persson, Gerhard Gröbner, Anders Sjöstedt, Marta Bally, Jonas Barandun, Bernt Eric Uhlin, Sun Nyunt Wai

## Abstract

Many pathogenic bacteria produce protein toxins that target and perturb host cell membranes. The secreted α-pore-forming toxins (α-PFTs) cause membrane damage via pore formation. This study demonstrates a remarkable, hitherto unknown mechanism by an α-PFT protein from *Vibrio cholerae*. As part of the MakA/B/E tripartite toxin, MakA is involved in membrane pore formation similar to other α-PFTs. In contrast, MakA protein alone induces tube-like structures in the acidic lysosomal host cell compartment. *In vitro* studies unravel the dynamics of tubular growth, which occur in a pH-, lipid- and concentration-dependent manner. A 3.7-Å cryo-electron microscopy structure of MakA filaments reveals a unique protein-lipid superstructure. In its active α-PFT conformation, MakA embeds its transmembrane helices into a thin annular lipid bilayer and spirals around a central cavity. Our study provides molecular insights into a novel tubulation mechanism of an α-PFT protein, revealing a new mode of action by a secreted bacterial toxin.

## Introduction

Several bacterial protein toxins are known to interact directly with target cell membranes by binding specific receptors, lipids or proteins on host cell membranes^1,2^. Both extracellular and intracellular bacterial pathogens produce and secrete host membrane-attacking pore-forming toxins (PFTs) to counteract host defenses, to promote colonization and spread, and to kill other bacteria^3,4^. PFTs can be categorized into two main groups, α-PFTs and β-PFTs, based on whether the secondary structure of the membrane-penetrating domain contains α-helices or β-barrels, respectively^3^. Generally, PFTs are secreted from bacteria in a monomeric, soluble form. Upon recognition and binding to the target cell membrane, the toxins undergo a conformational change, interact with the membrane, dimerize, and oligomerize, leading to the formation of membrane pores^5^.

*Vibrio cholerae,* a Gram-negative bacterium, is the causal organism of the diarrheal disease cholera^6^. Cholera toxin (CT) and toxin co-regulated pilus (TCP) are the main virulence factors of *V. cholerae* that cause disease in mammalian hosts^7,8^. Most environmental *V. cholerae* isolates do not produce CT and TCP^9^. Nevertheless, these bacteria are considered pathogenic since they have been associated with secretory diarrhea and may cause wound infections and sepsis^10^. *V. cholerae* strains lacking the cholera toxin-encoding genes often contain a set of genes coding for other secreted virulence factors, including hemolysin, hemagglutinin protease, RTX toxin, and multiple lipases that jointly may play a role in pathogenesis^10^.

Using *Caenorhabditis elegans* and *Danio rerio* (zebrafish) as host models for bacterial predatory interactions and infection in aqueous environments, respectively, we obtained evidence for a new *V. cholerae* cytotoxin denoted MakA (motility-associated killing factor A)^11^. The *V. cholerae* gene *makA* (*vca0883)* is localized in a gene cluster together with *makC* (*vca0881*), *makD* (*vca0880*)*, makB* (*vca0882*) and *makE* (*vca0884*). This gene cluster has been found in different *V. cholerae* strains, including CT-negative isolates^11,12,13^. The crystal structure of MakA revealed that it belongs to the ClyA α-PFT family^11^, named after the potent, one-component, pore-forming toxin ClyA which is expressed from a monocistronic operon in *E. coli*^14,15^. Our recent studies of the proteins encoded by the *mak* genes in *V. cholerae* demonstrated that MakA can form a tripartite cytolytic complex with MakB and MakE, whereas neither of the three proteins display cytolytic activity on their own^13^. Other family members of α-PFT are found among the bipartite toxins from *Yersinia enterocolitica* (YaxAB)^16,17^ and *Xenorhabdus nematophila* (XaxAB)^17^, as well as among the tripartite toxins from *Bacillus cereus* (NheABC and HblL_1_L_2_B)^18^, *Aeromonas hydrophila* (AhlABC)^19^ and *Serratia marcescens* (SmhABC)^20^. The bipartite and tripartite PFTs require the combined action of all protein partners to induce pore formation in the target membranes, and there is evidence that protein interactions occurs in a specific order for maximum cytolytic activity^13,19,20^. However, there are still questions about how many molecules of each subunit protein are required to form a pore, how they interact with each other, how structural conformational changes occur, and how protein moieties are involved in the interaction with the host membrane. In addition, it remains possible that some of the subunit proteins can be separately released from the bacteria and thereby exhibit biological effects on their own. Secretion of the MakA/B/E proteins from *V. cholerae* was shown to be facilitated via the bacterial flagellum^11,13^. However, about 10% of MakA was secreted from *V. cholerae* lacking the flagellum suggesting an alternative route of secretion, in contrast to MakB and MakE, which displayed a more definitive flagellum-dependent secretion^11,13^.

Our earlier studies with MakA and cultured mammalian cells showed that the protein binds to the target cell membrane and, upon internalization, may accumulate in the endolysosomal membrane, causing lysosomal dysfunction, induction of autophagy and apoptotic cell death^21,22^. Moreover, MakA can modulate host cell autophagy in a pH-dependent manner^23^.

The present study aimed to further characterize the mechanism(s) behind MakA-induced lysosomal membrane tubulation and the pH-dependent molecular interaction(s) between MakA and host cell membranes. We found that under low pH conditions, MakA bound to purified lysosomes and to liposomes prepared from total epithelial cell lipid extracts (ECLE). The insertion of MakA into lysosomes and ECLE liposomes resulted in the formation of tubular assemblies. Cryo-electron microscopy (cryo-EM) analysis of these assemblies revealed an unusual helical structure formed by MakA and lipid spirals. The observed structure revealed that MakA monomers adopted conformational arrangements typical of active membrane-bound α-PFTs while they assembled into an atypical polymeric superstructure. Large structural rearrangements, presumably induced by the lowered pH, were necessary for the transition from the inactive soluble form to the extended active toxin form. Interaction of these MakA structures with cell membranes could lead to cell death in the *in vitro* setting. MakA is the first *V. cholerae* protein that engages target membranes to form nanotubes by polymerizing as a helical structure together with a lipid spiral.

## Results

### pH-dependent formation of tubular structures in lysosomes and on cell membranes by MakA

In recent *in vitro* studies with epithelial cells, we found that internalized MakA protein accumulated in the acidic endolysosomal compartment where it caused formation of tube-like structures and induced lysosomal permeability^22^. Our findings prompted us to examine how the observed lysosomal tubulation and lysosomal dysfunction might be caused by MakA. Upon treatment of Caco-2 cells with MakA (250 nM, 18 h), we observed co-localization of the Alexa568-labeled MakA (Alexa568-MakA) with GFP-LAMP1 or lysotracker in tubular structures (**Fig. 1a** **and Supplementary Fig. 1a**). The tubulation in the lysosomes was further confirmed by transmission electron microscopy (TEM) of lysosomes isolated from MakA-treated HCT8 cells (**Fig. 1b**). To investigate if MakA can also induce tubulation of lysosomes outside the intracellular environment, we purified lysosomes from untreated HCT8 cells and exposed them to native MakA or to Alexa568-MakA. Both confocal microscopy and TEM analysis revealed aggregation and tubulation of lysosomes at pH 5.0 (**Fig. 1c-d**). In addition, a majority of the lysosomes showed well-organized tubulation when exposed to MakA at pH 6.5 (**Fig. 1d**). In contrast, we did not observe any MakA-induced tubular structures in lysosomes at pH 7.0 (**Fig. 1d**). Western blot analysis confirmed pH-dependent binding of MakA to lysosomes (**Fig. 1e**). Alexa568-MakA was subsequently shown to bind epithelial cells in a pH-dependent manner (**Fig. 1f-g**). To determine the kinetics of MakA binding to the target cells, HCT8 cells were exposed to Alexa568-MakA at pH 5.0, and live-cell imaging was performed using spinning disc confocal microscopic analysis. Consistent with our earlier findings^21^, we observed accumulation of Alexa568-MakA on the plasma membrane, including filipodia-rich tubular structures, in a time-dependent manner (**Fig. 1h** **and Supplementary Fig. 1b-c)**. The time scale of MakA binding to individual HCT8 cells ranged from ∼40 minutes to 4 hours after Alexa568-MakA treatment (**Fig. 1f-h** **and Supplementary Fig. 1b-c**). Ultimately, Alexa568-MakA was detected on the plasma membrane of the entire cell population, with most cells positive for tubular structures protruding out from the plasma membrane (**Fig. 1f and 1h**). Taken together, these results suggest that MakA can cause tubulation of both endolysosomal membranes and plasma membranes in a pH-dependent manner.

**Fig. 1:**
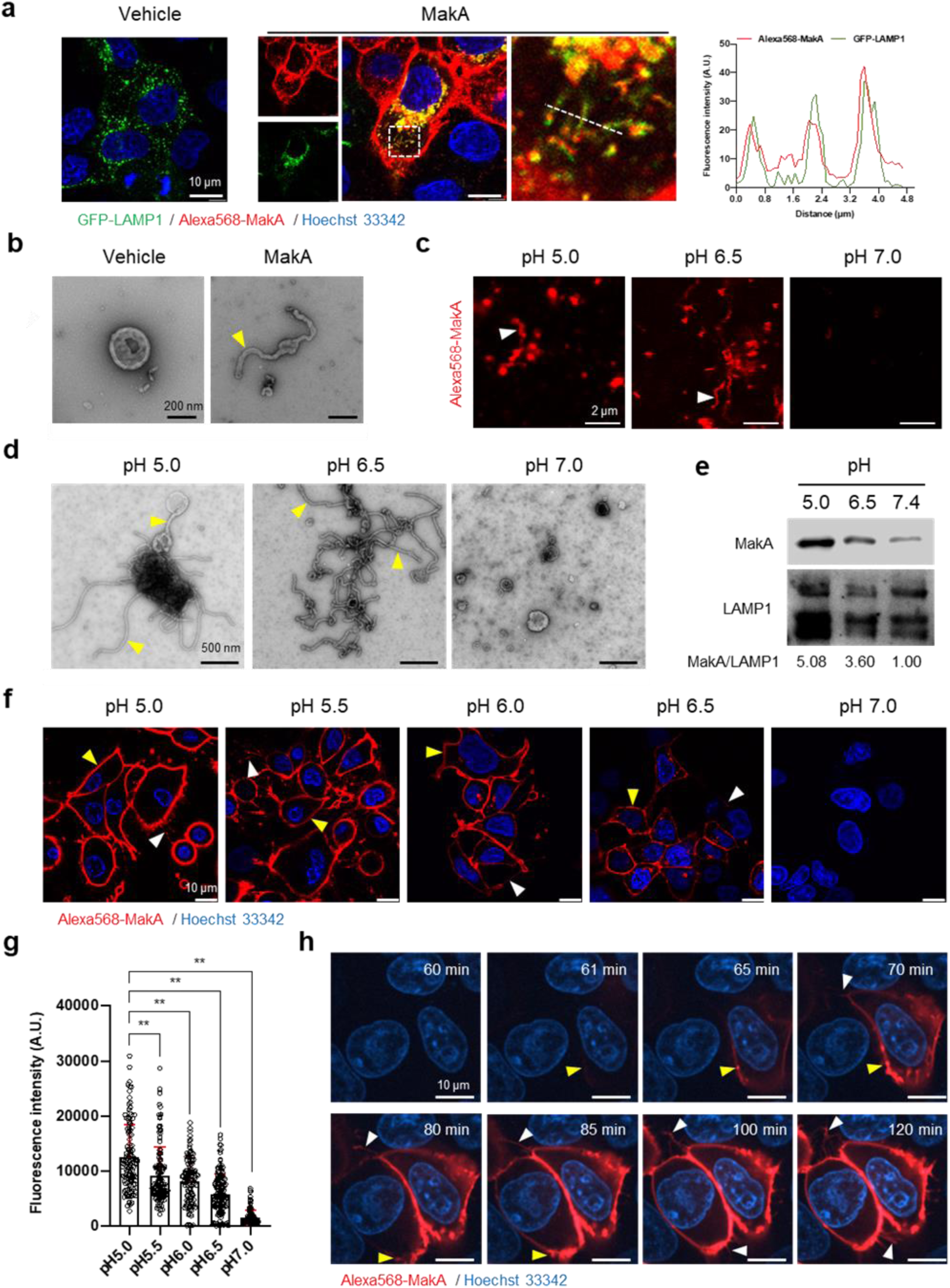
pH-dependent tubulation of lysosomes and binding to epithelial cells by MakA. **a** Caco-2 cells transfected with GFP-LAMP1 were treated with vehicle or Alexa568-MakA (250 nM, 18 h). Nuclei were counterstained with Hoechst 33342. The line graph to the right indicates the accumulation of Alexa568-MakA in GFP-LAMP1-positive tubular lysosomes. Pearson correlation co-efficient was used to calculate Alexa568-MakA (red) co-localization with GFP-LAMP1 (green) along with the tubular structures. Scale bars, 10 μm. **b** Representative electron micrographs of lysosomes purified from vehicle and MakA (250 nM, 24 h) treated HCT8 cells. Scale bars, 200 nm. The yellow arrowhead indicates a tubular structure found with lysosomes from MakA-treated cells. **c** The pH-dependent binding of Alexa568-MakA to purified lysosomes isolated from HCT8 cells: White arrowheads point to the tubular structures observed with MakA-treated purified lysosomes. Images shown for specimens from different pH conditions were acquired using the same settings of the microscope. Scale bars, 2 µm**. d** Representative electron micrographs of purified lysosomes treated with MakA (1 µM) under different pH conditions. Yellow arrowheads indicate tubular structures appearing at low pH. Scale bars, 500 nm. **e** Western blot analysis of samples from lysosome pull-down assays performed with lysosomes treated with MakA (250 nM, 60 min) under different pH conditions. Lysosome-bound MakA was detected with anti-MakA antiserum. Immunodetection of LAMP1 was used as a reference and the MakA/LAMP1 ratio was determined for the quantification of relative MakA amounts. **f** HCT8 cells were exposed to Alexa568-MakA (500 nM, 4 h) under different pH conditions and visualized live under by confocal microscopy. Nuclei were counterstained with Hoechst 33342 (blue). Yellow arrowheads indicate cell membrane association, while white arrowheads indicate MakA-positive tubular structures. The different images were acquired using the same microscope settings. Scale bars, 10 μm. **g** The histogram indicates quantification of cell-bound Alexa568-MakA (n = 100 cells) as shown in (f). Data from two independent experiments are presented as mean ± s.e.m.; one-way analysis of variance (ANOVA) with Dunnett’s multiple comparisons test. **p≤0.01. **h** Still images of HCT8 cells exposed to Alexa568-MakA (500 nM) at pH 5.0. Yellow arrowheads indicate the initial binding site of MakA and white arrowheads indicate the appearance of MakA-positive tubular structures in a time-dependent manner. Nuclei were counterstained with Hoechst 33342. Scale bars, 10 μm. **Source data for** figure 1e; Uncropped MakA-detected membrane.

### pH-dependent epithelial cell toxicity and formation of tubular structures on erythrocytes by MakA

To further assess the effect of pH on the binding of MakA to HCT8 epithelial cells, we performed Western blot analysis using anti-MakA antiserum (**Fig. 2a**). We observed that MakA bound to the cells in a pH-dependent manner, and moreover, that there seemed to be a pH-dependent formation of stable MakA oligomers bound to the epithelial cells. To determine if MakA binding and oligomerization at the target cell membrane correlated with cytotoxic effect, HCT8 cells were exposed to MakA, and cell toxicity was quantified by a propidium iodide uptake assay using flow cytometry and fluorescence microscopy (**Fig. 2b** **and Supplementary Fig. 2a**). In addition to causing pH-dependent toxicity of HCT8 cells, MakA was similarly toxic to other colon cancer cells, Caco-2 and HCT116 cells, as assessed by the MTS cell viability assay (**Supplementary Fig. 2b**).

**Fig. 2:**
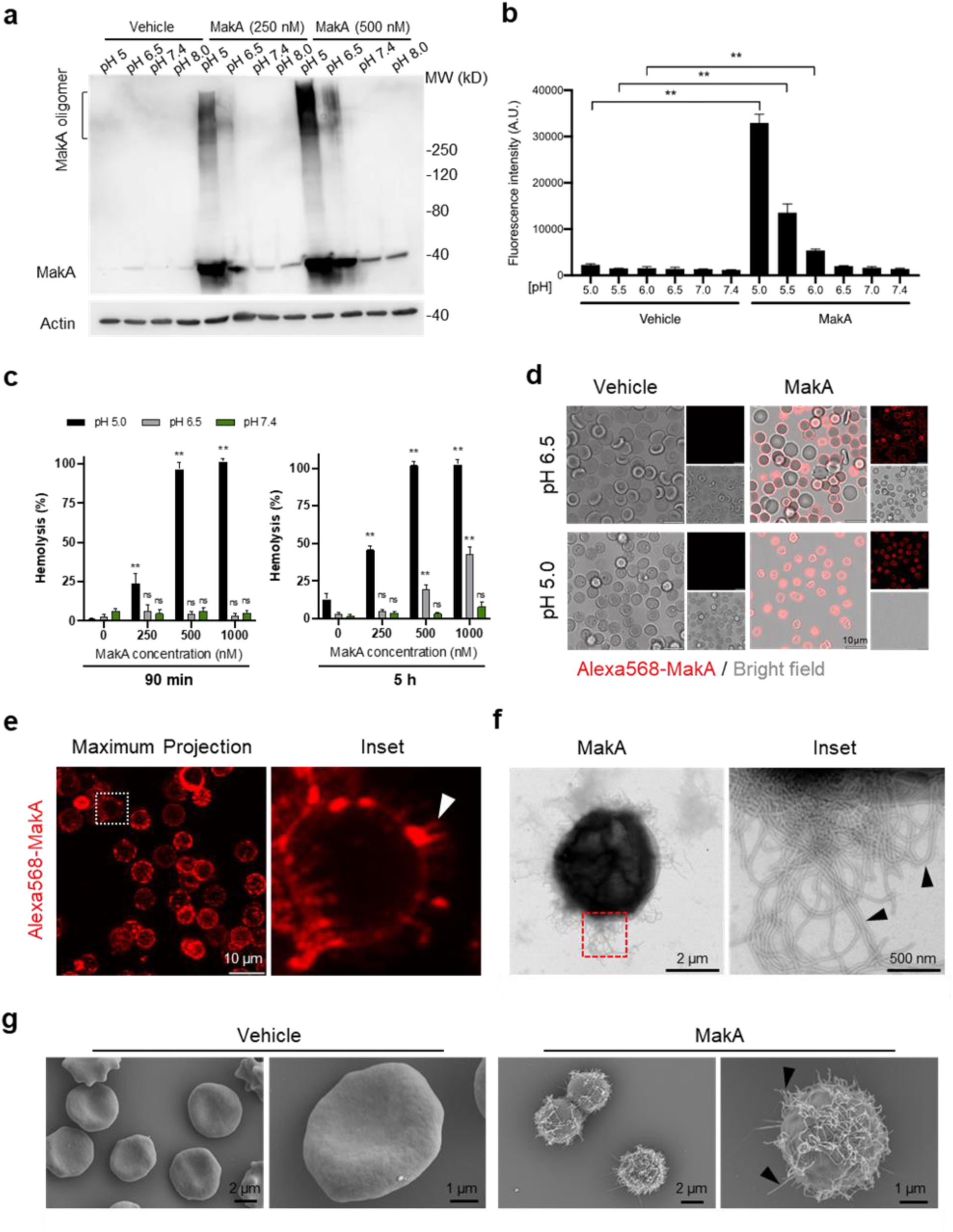
pH-dependent MakA binding to target cell membranes and induction of tubulation on erythrocytes. **a** Western blot analysis of HCT8 cells treated with increasing concentrations of MakA under different pH conditions for 4 h. Data are representative of two independent experiments. Cell-bound MakA was detected with MakA-specific antibodies, and actin was used as a loading control. **b** HCT8 cells were treated with MakA (500 nM, 4 h) under different pH conditions, and cell toxicity was monitored by assaying the uptake of propidium iodide. Fluorescence values for propidium iodide were recorded by flow cytometry. Data are representative of three independent experiments; bar graphs show the mean ± s.d. Significance was determined from biological replicates using a non-parametric t-test. **p≤0.01. **c** Human erythrocytes suspended in citrate buffer of different pHs were exposed to increasing concentrations of MakA for 90 min (left panel) and 5 h (right panel). MakA-induced hemolysis of erythrocytes was normalized against erythrocytes treated with Triton X-100 (0.1%), and data was expressed as a percentage (%). Data are representative of six readouts from two independent experiments; bar graphs show mean ± s.d. Significance was determined from biological replicates using a non-parametric t-test. **p≤0.01, *p≤0.05, or ns = not significant. **d** Human erythrocytes (0.25%) in phosphate-buffered saline (PBS) were allowed to adhere to the glass surface for 10 h, followed by buffer exchange to citrate buffer (pH 5.0 or pH 6.5). The erythrocytes were treated with Alexa568-MakA (500 nM, 3 h), and cell-bound MakA was detected by confocal microscopy. Scale bars, 10 µm. **e** The image shows a maximum z-stack projection of the human erythrocytes treated with Alexa568-MakA (pH 6.5 in citrate buffer). The white arrowhead in the right panel indicates the accumulation of Alexa568-MakA in tubular structures at the surface of erythrocytes. Scale bars, 10 µm. **f** Transmission electron microscopy (TEM) images of erythrocytes treated with vehicle or MakA (500 nM) for 90 min and stained with 1.5% uranyl acetate solution. Black arrowheads in the enlarged part of the image to the right indicate the presence of tubular structures present on the surface of the liposome. **g** Scanning electron microscopy (SEM) images of erythrocytes treated with MakA (500 nM, 90 min) in citrate buffer (pH 6.5). Representative examples of imaged erythrocytes indicate that the formation of tubular structures occurred throughout the surface of MakA treated erythrocytes. Scale bars, 2 µm. Figure 2**; source data** Uncropped western blot membranes

Erythrocytes are widely used as a cell model to investigate the cytolytic activity of the toxins that belong to the ClyA pore-forming toxins family^14,17,18,24,25^. To test if pH-dependent binding and oligomerization of MakA may cause hemolysis of erythrocytes, human erythrocytes were exposed to increasing concentrations of MakA at different pH conditions for either 90 min or 5 h (**Fig. 2c**). When erythrocytes were exposed to MakA at pH 5.0, hemolysis was observed in a concentration dependent manner within 90 min (**Fig. 2c**). In contrast, MakA failed to induce hemolysis of erythrocytes at pH 6.5 or pH 7.4 during the 90 min treatment. A detectable, but low level of hemolysis was observed after 5 h with erythrocytes exposed to MakA at pH 6.5 (**Fig. 2c**). With confocal microscopy, we observed pH-dependent binding of Alexa568-MakA to erythrocytes. A majority of erythrocytes at pH 5.0 and 6.5 were covered by Alexa568-MakA whereas there was virtually no MakA binding observed at pH 7.4 (**Fig. 2d** **and Supplementary Fig. 2c**). Notably, we detected the presence of tubular structures in association with MakA at the red blood cell surface, as shown by a maximum 3D projection of the z-stack images of erythrocytes (**Fig. 2e**). The presence of MakA-induced tubular structures on the surface of erythrocytes was further observed by TEM and scanning electron microscopy (SEM) (**Fig. 2f-g**). Together, these results suggest that MakA could accumulate in a pH-dependent manner at the surface of both epithelial cells and erythrocytes, thereby inducing formation of tubular structures that ultimately might lead to cell lysis.

### MakA induction of tubular structures on liposomes lacking other proteins

To investigate if tubulation of host membranes in response to MakA would require a specific membrane protein or receptor, we created protein- and cytosol-free liposomes using an epithelial cell total lipid extract (ECLE) isolated from HCT8 cells. After addition of MakA, the liposomes were pelleted by centrifugation and the presence of MakA in either the pellet or the supernatant was detected by Western blot analysis (**Supplementary Fig. 3a**). The results indicated that more MakA was associated with the liposomes at pH 5.0 and 6.5 than at pH 7.4. The interaction between MakA and ECLE liposomes at pH 6.5 was quantified by surface plasmon resonance (SPR) analysis (**Fig. 3a**). MakA displayed significant interaction with ECLE liposomes, with an estimated K_D_ of 49.2 nM. Importantly, MakA at the highest tested concentration (200 nM) failed to interact with the liposomes prepared from zwitterionic 1-palmitoyl-2- oleoyl-sn-glycero-3-phosphocholine (POPC), used as a negative control (**Fig. 3a**).

**Fig. 3:**
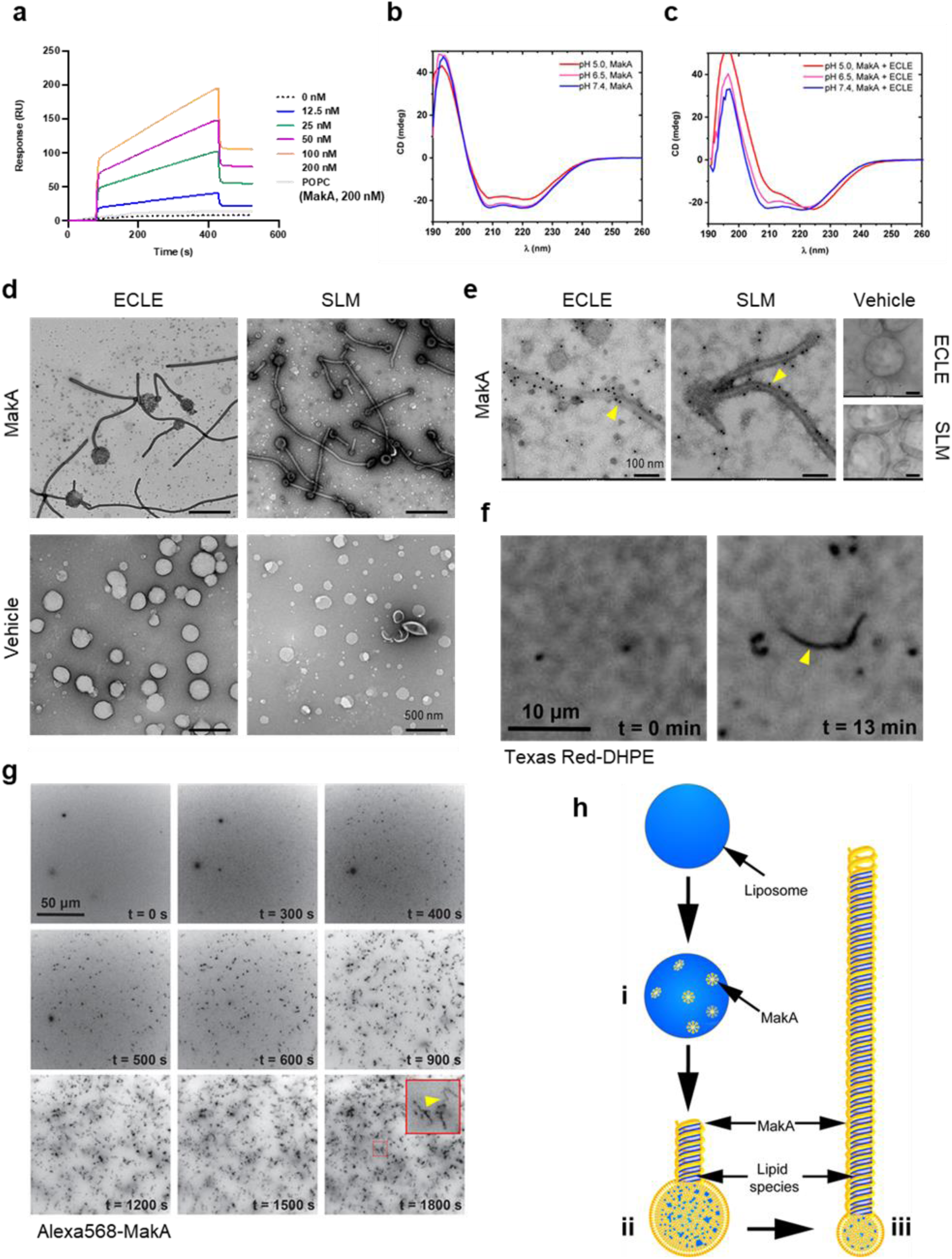
pH-dependent formation of protein-lipid tubular structures from MakA interaction with liposomes. **a** Surface plasmon resonance (SPR) assay showing direct binding of MakA (0 to 200 nM) to ECLE liposomes (120 mM citrate buffer, pH 6.5). The liposomes were immobilized on an SPR sensor chip L1 (dotted line with 0 nM is buffer control and grey line with immobilized POPC was used as a negative control). Control flow cell background was subtracted from the experimental cell before final data processing. The K_D_ values (49.2 nM) were calculated by using the BioLogic scrubber 2 software. **b** Far-UV CD spectra of native MakA of MakA bound to ECLE liposomes under different pH condition. CD spectra were recorded in 5 mM citrate buffer using 3 μM MakA protein. The absorption intensity measured from the control solution, containing buffer only, was subtracted to account for background absorption. **d** ECLE or synthetic lipid mixture (SLM) liposomes were treated at pH 6.5 with vehicle (Tris 20 mM) or MakA (3 µM) for 90 min and stained with 1.5% uranyl acetate solution. Images were captured with transmission electron microscopy (TEM). White arrowheads indicate tubular structures and blue arrowheads indicate MakA oligomeric structures present in the background of liposomes. Scale bars, 200 nm. **e** The ECLE or SLM liposomes were treated with vehicle (Tris 20 mM) or MakA (3 µM) for 90 min and stained with 1.5% uranyl acetate solution. MakA was detected with anti-MakA antibodies, followed by immunogold labeling and imaging by TEM. Scale bars, 200 nm. **f** Selected inverted grayscale images from time-lapse epifluorescence microscopy [Movie 1] obtained after incubating SLBs (prepared from SLM+TxRed liposomes) with MakA (3 μM) at pH 6.5. The fluctuating tubules (yellow arrowhead) are visible due to their TxRed-DHPE lipid content. Scale bar, 10 μm. **g** Selected inverted grayscale images from time-lapse epifluorescence microscopy [Movie 2] obtained after incubating SLBs (prepared from SLM liposomes) with Alexa568-MakA (3 μM) at pH 6.5. Panels illustrate key steps during the transformation of SLBs into fluctuating tubules. Arrowhead (yellow) in the inset indicate appearance of a Alexa568-MakA positive tubular structure. Scale bar, 50 μm. **h** Schematic representation of how MakA insertion into the liposome can result in formation of a MakA- and lipid-positive tubular structure under low pH (6.5 or lower) conditions. Upon addition of MakA (yellow) the liposome surface (blue) will become covered by the protein (steps i and ii) and the MakA oligomerisation initiates formation of a tube-like structure that gradually appears to deplete the liposome of lipids when the protein-lipid tube assembly continues to grow in length (step iii).

To determine the effect of pH on MakA’s conformational properties, the protein was subjected to circular dichroism (CD) spectroscopy analysis at different pH in the absence or presence of ECLE liposomes (**Fig. 3b-c**). In the absence of liposomes, the CD spectra indicated a decrease in the α-helical content of the protein when in the acidic environment. This decrease in α-helical content of MakA was restored when ECLE liposomes were present, suggesting that liposomes somehow stabilized the structure of MakA (**Fig. 3b-c** **and Supplementary Table 1**). TEM analysis of MakA at different pH indicated that it formed oligomeric structures in a pH-dependent manner (**Supplementary Fig. 3b**).

Further examination by TEM addressed if there were morphological changes in the ECLE liposomes upon exposure to increasing concentrations of MakA at pH 6.5 (**Fig. 3d** **and Supplementary Fig. 3c**). Similar to the tubulation observed for the target cell membranes and lysosomes, MakA triggered tubulation of the ECLE liposomes in a concentration-dependent manner (**Supplementary Fig. 3c**). Concomitant with the assembly of tubular structures emanating from the liposomes, the size of the liposomes appeared to shrink as the tubules grew up to several micrometers in length. Ultimately, the entire liposome seemed to be transformed into long tubules (**Fig. 3h** **and Supplementary Fig. 3c-e**). The tubulation of ECLE liposomes was also observed by confocal microscopy upon treatment with Alexa568-labeled MakA (**Supplementary Fig. 3d-e**). In the same population of small liposome particles, we also detected Alexa568-MakA-positive large lipid vesicles (5-10 μm in size, less than 1% of the entire liposome fraction). The z-stack projection suggested that the whole lipid vesicle was decorated with a bundle of fluctuating tubules (**Supplementary Fig. 3e**). To further assess whether or not any protein or glycolipid receptor mediated the observed membrane tubulation by MakA, liposomes were prepared from a well-defined synthetic lipid mixture (SLM); whose composition was inspired by the distribution of lipids found in the plasma membrane of HeLa cells^26^. Tubulation of the SLM liposomes by MakA was observed by TEM (**Fig. 3d**). In addition to the tubular structures, we observed a large number of well-organized, star-shaped oligomeric particles of MakA among the ECLE liposomes (**Supplementary Fig. 3c**). Furthermore, the presence of MakA protein in the tubular structures was evidenced by immunogold staining using MakA-specific antibodies (**Fig. 3e**). By fluorescence microscopy we were able to visualize tube growth originating from a supported lipid bilayer (SLB) prepared from SLM liposomes containing the fluorescent lipid Texas Red-DHPE, demonstrating that the tubes also contain lipids from the SLB (**Fig. 3f** **and Movie 1**).

We next investigated the kinetics of MakA protein-lipid tubulation. Using fluorescence microscopy, we found that administering Alexa568-MakA to a SLB prepared from SLM liposomes prompted a significant and highly dynamic membrane remodeling (**Fig. 3g** **and Movie 2**). Within 10 minutes, Alexa568-MakA binding to the SLBs resulted in formation of MakA-associated tubules of various sizes (**Fig. 3g**). Based on these findings, we propose a schematic model for how the MakA-liposome interaction can result in the formation of the observed protein-lipid tubular structures (**Fig. 3h**). At pH 6.5 or lower, MakA may adopt a conformation that allows the protein to insert into the lipid membrane in the form of an oligomer assembly that can start to spiral around the lipids of the membrane leading to formation of a growing tube structure. Concomitantly the size of the vesicle appear to shrink, and the tube may grow up to several micrometers in length. Our results suggest that the MakA-lipid tubulation can occur without the involvement of other proteins or some specific protein receptor under the pH conditions tested.

### Structure of the MakA filament

We used helical reconstruction to solve the cryo-EM structure of a MakA filament assembled *in vitro* in the presence of ECLE liposomes at pH 6.5 and high protein concentrations (**Fig. 4a** **and Supplementary Fig. 4**). An initial 2D classification allowed us to identify repetitive elements and measure a helical repeat distance of ∼216 Å (**Fig. 4b** **and Supplementary Fig. 5a**). A subsequent investigation of the layer line distances in a collapsed power spectrum of selected well-resolved class averages confirmed this distance (**Supplementary Fig. 5b**). Next, we performed a preliminary 3D refinement of filament segments from well-defined 2D class averages without imposing symmetry (**Supplementary Fig. 4**). This volume was visually inspected to deduce the helical symmetry parameters (**Supplementary Fig. 5c-d**). One repeating element of the right-handed spiral consists of 37 tetramers that complete five turns around the helical axis, spanning a length of 216.5 Å and a diameter of 322 Å. This arrangement results in an axial rise of 5.85 Å per subunit and a helical twist of 48.65° (**Fig. 4b** **and Supplementary Fig. 5d**). Application of these initially calculated values with local searches in a 3D refinement further optimized symmetry parameters, resulted in a cryo-EM map at an overall resolution of 3.7 Å (**Supplementary Table 1 and Supplementary Fig. 6**). The obtained cryo-EM map features a well-resolved central transmembrane helix (TMH) region and a less well-resolved peripheral region (**Fig. 4c**). We isolated two MakA tetramers from the segments using signal subtraction and subjected the resulting particles to 3D classification and refinement to improve the peripheral density and connectivity. The clear connectivity of the obtained density map (**Fig. 4d** **and Supplementary Fig. 6e**) allowed for reliable secondary structure placement using the MakA soluble state crystal structure (PDB-6EZV^11^). High-resolution features in the helical reconstruction (**Fig. 4c** **and Supplementary Fig. 6f**) allowed for *de novo* model building of structural elements in the central region. However, due to continuous rotation along the filament axis, flexibility (**Supplementary Fig. 4**), and the conformational difference with respect to the crystal structure, the MakA tail domain structure is less reliable and modeled with poly-alanine secondary structure elements.

**Fig. 4:**
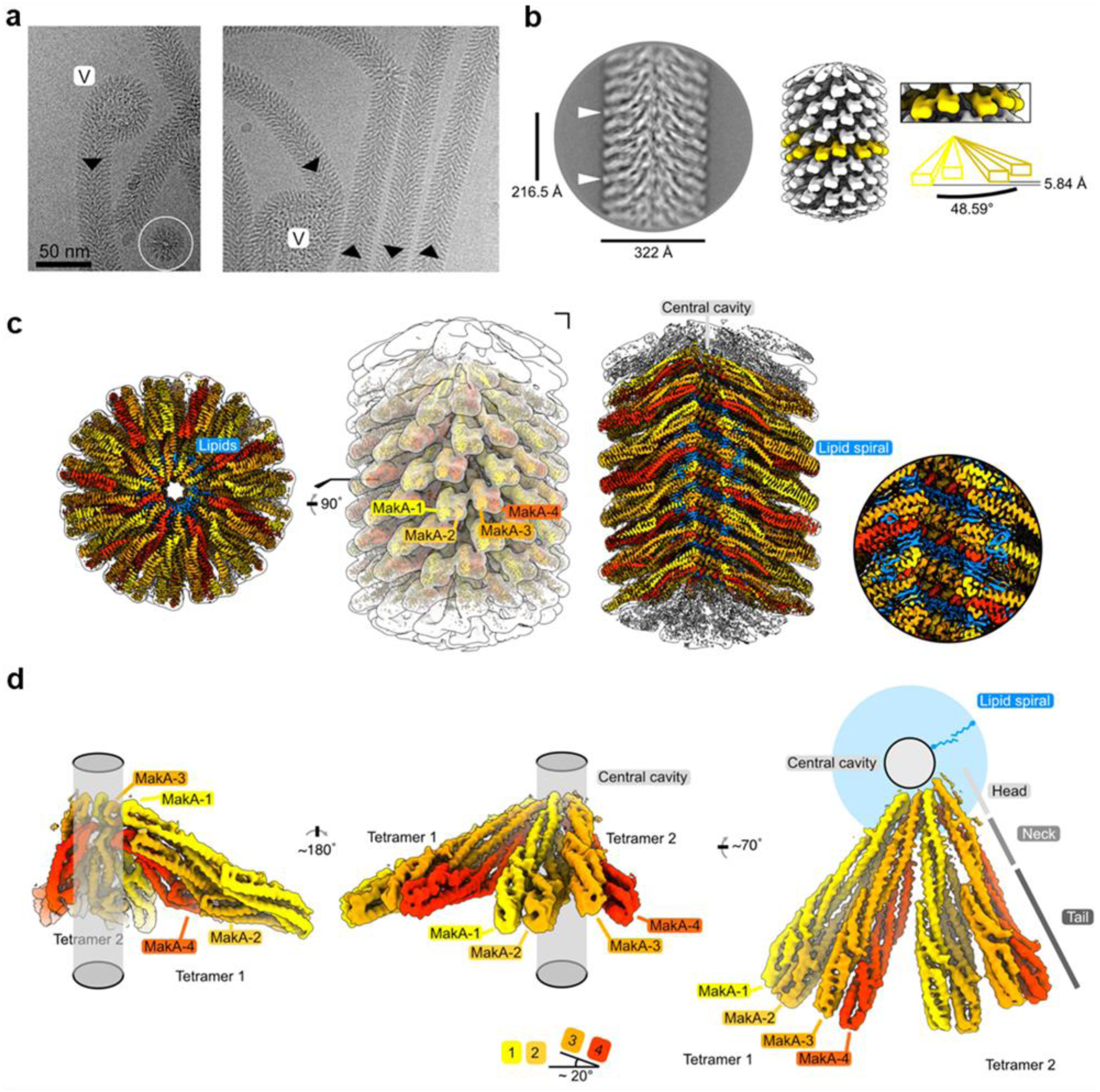
Cryo-EM structure of the membrane-bound MakA filament. **a** Representative cryo-EM micrograph sections showing MakA filaments emerging from or ending in a membranous vesicle (V; vesicle). The black arrows indicate the directionality of the filaments. A top-view of the filamentous tube is visible in the first micrograph within the white circle. **b** A 2D class average with filament diameter indicated below and the repeat distance labeled on the side. An example of a visually repeating element is indicated with white arrows. The right side depicts a low-pass filtered cryo-EM volume with eight repeating subunits colored in gold next to a zoom-in of two blades. A schematic representation of the two repeating units is visualized underneath the zoom-in with a helical twist (48.59°), and rise (5.84 Å) indicated. **c** Overall cryo-EM volume (EMD-13185) and slab views of the MakA filament superimposed onto a semi-transparent, white, 20-Å, low-pass filtered map. The four MakA subunits, belonging to one tetramer, are colored in shades of gold and orange-red and labeled. The different densities between the protein blades belonging to a lipid bilayer is colored in blue. **d** Rotationally related views of the signal of subtracted and focused-refined cryo-EM volume of two tetramers (EMD-13185-additional map 1) are shown with a schematic representation of the central cavity (transparent grey) and the lipid spiral (blue). Common structural elements of the alpha-cytolysin family protein-fold are indicated in grey (Head, Neck, and Tail). The 20°-rotation of the tail domain between two dimers within the asymmetric unit is shown schematically below the central panel.

MakA oligomerizes into a filamentous structure growing from or ending in membranous vesicles (**Fig. 4a, Supplementary Fig. 4**). The building blocks of this filament are formed by two MakA dimers (**Fig. 4b and 4d**) that organize into a pinecone-like architecture, spiraling around a central cavity (**Fig. 4c**). From the top view along the filament axis, the helix features a propeller-like structure with a weak, annular density embedded in between the blades formed by MakA (**Fig. 4c**). This density resembles lipid tails and contains some spherical features, which could be associated with phospholipid heads, suggesting the presence of a thin phospholipid bilayer that spirals around the central cavity of the filament (**Fig. 4c**). Interestingly, the annular density is located between the transmembrane helices of MakA (**Fig. 4d**), indicating that the active toxin form interacts with lipid vesicles and starts to oligomerize by internalizing membrane lipids.

### A significant conformational change is required to adopt the membrane-bound state

The basic building block of the observed protein-lipid filament is formed by four MakA subunits in a membrane-bound active conformation (**Fig. 5a,b**). This conformation of MakA is significantly different from the previously reported soluble state structures resembling the inactive form (PDB-6DFP and PDB-6EZV^11^). In the soluble form, a C-terminal tail (res. 351-365, **Fig. 5c**, purple) inactivates the predicted transmembrane domains by forming a β-tongue, consisting of three β-sheets^11^, that shields the hydrophobic residues from the surrounding solvent. This shielding characteristic is well-described for the soluble form of the ClyA pore-forming toxin family, including ClyA, Hbl-B, NheA, and AhlB^17,19, 27–30^. MakA undergoes a structural change comparable to the opening of a Swiss army knife blade when it shifts from a soluble inactive to a membrane-bound active state, where the helix bundle of the tail region represents the handle, the transition from the tail to the neck region forms two hinges, and the β-tongue together with 4 and 5 resembles the blade that folds out (**Fig. 5c**, α4 & α5; light & dark green). Additionally, the β-tongue changes its secondary structure and, together with α4 and α5/α6, forms two extended helices (**Fig. 5c**). This significant conformational change leads to the formation of two TMHs and a short loop region (res. 219-221) between α4 and α5. Interestingly, despite certain similarities, all four copies of MakA adopt different conformations within the tetramer, indicated by MakA-1 to MakA-4 (**Fig. 4** **and** **Fig. 5**). The most significant structural differences between the four MakA conformations can be observed in the neck and head domains, connected via a hinge with the tail domains (**Fig. 5c**). The hinge allows for different degrees of bending (opening up of the Swiss army knife), while the neck helices α4 and α5 and the loop region (res. 219-221) display a high degree of plasticity. For the peripheral tail domain, two major conformations can be observed. In MakA-2 and MakA-4, this region superimposes well with the crystal structure, whereas helix α6 of MakA-1 and MakA-3 moves by almost 10 Å to extend the length of α5. While the N-terminal helices of the stretched MakA states are reduced compared to those of the kinked MakA forms, the C-terminal β-strand (res. 351-365), covering the TMH in the soluble state, is disordered in all four subunits.

**Fig. 5:**
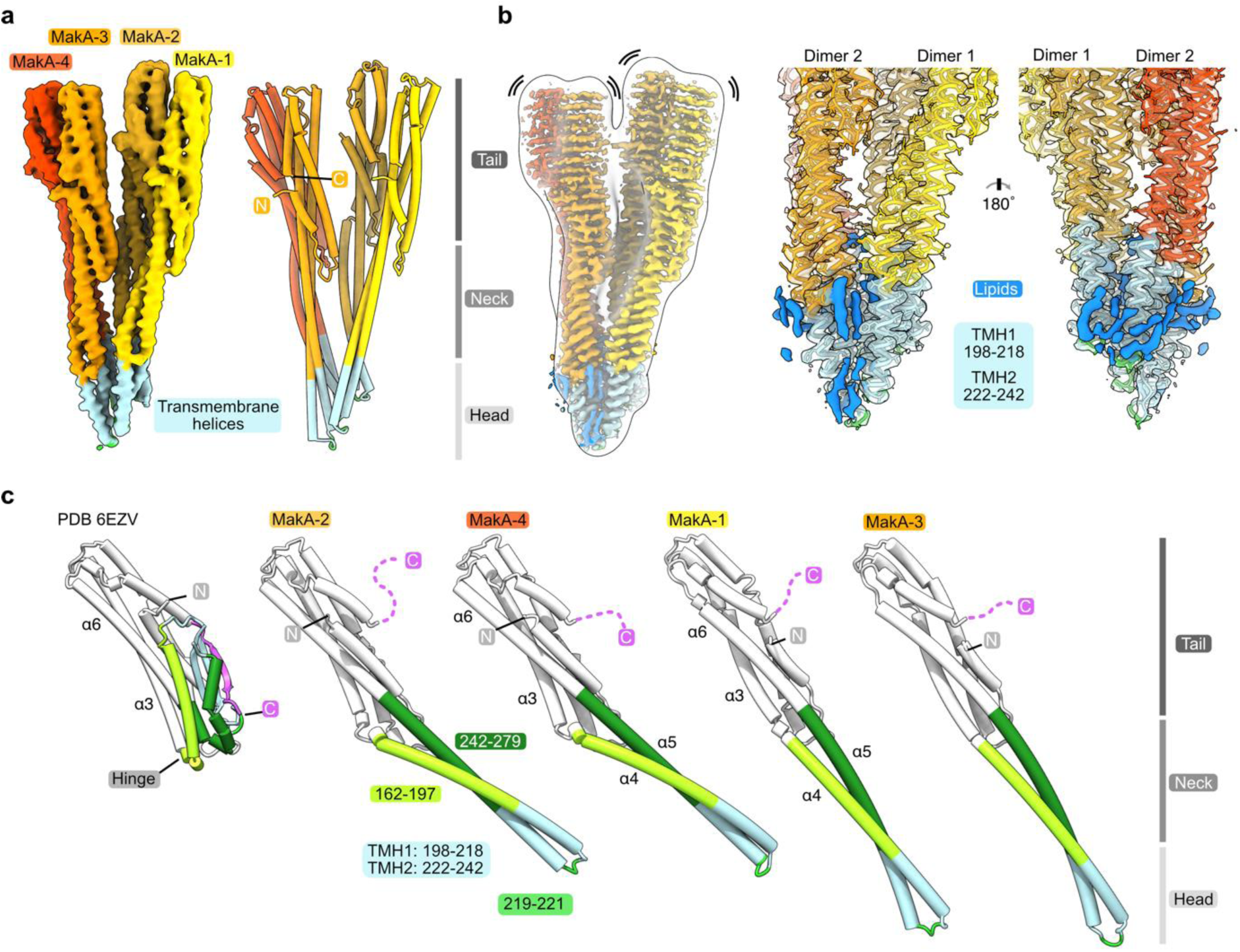
Conformational plasticity of MakA in the membrane-bound filamentous state. **a** The cryo-EM density of a MakA tetramer (EMD-13185-additional map 1) is shown next to a structural model. The individual protein domains (Head, Neck and Tail) and the visible N- and C-terminal are indicated. **b** The cryo-EM density of a MakA tetramer, obtained by helical reconstruction (EMD-13185), is shown in isolation, colored and oriented as the structural model in (**a**). The cryo-EM volume is superimposed with a white, transparent, 20-Å, low-pass filtered volume. Additional density areas in the transmembrane helix (TMH) region, presumably belonging to lipids, are colored in blue. **c** The crystal structure of monomeric MakA (PDB-6EVZ^11^) with retracted neck and head domain is shown next to the four individual subunits of the membrane-bound state of MakA in cartoon representation (PDB-7P3R). All structural models were superimposed based on the tail region (in white) and displayed in the same orientation next to each other with increasing length depicting flexing of the neck and head domain as well as the TMH.

## Discussion

Recent studies have revealed how different bacterial species, notably *B. cereus*, *A. hydrophila*, *S. marcescens* and *V. cholerae*, have the capability to produce structurally similar tripartite protein complexes that assemble on host cell membranes as pore structures that are cytolytic^13,18–20^. In the case of *V. cholerae,* it has become evident that the MakA component of the tripartite complex, when presented alone to mammalian cells, is effectively internalized and accumulates on endolysosomal membranes, leading to induction of autophagy and apoptotic cell death^21–23^. Herein, we show that MakA is able to produce a novel protein-lipid polymeric superstructure at low pH (6.5 or below) that perturbs host cell membranes.

Several prokaryotic proteins are known to polymerize in the presence of a matrix such as a lipid membrane or DNA. This type of assembly is referred to as a collaborative filament^31^. In the presence of lipid membranes, collaborative filaments are assembled by a bridging protein and lipid on the membrane in a sequence-specific manner and by sensing membrane curvature^31^. It was demonstrated that membrane-mediated clustering of Shiga toxin molecules and the formation of tubular membrane invaginations are essential steps in the clathrin-independent Shiga toxin uptake process^32^. In earlier studies, pH-dependent membrane insertion and increased cytotoxic activity were demonstrated for different bacterial toxins, including anthrax toxin from *Bacillus anthracis*^33^, diphtheria toxin from *Corynebacterium diphtheriae*^30^, VacA from *Helicobacter pylori*^33^, perfringolysin from *Clostridium perfringens*^34^, and listeriolysin O from *Listeria monocytogenes*^35^. Recent research on the pre- and pore forms of mammalian pore-forming toxin, mammalian perforin 2 (mPFN2), provide interesting insights into the pore generation process. It was demonstrated that pre-pore-to-pore transformation occurs at an acidic pH, which is accomplished by a 180° rotation of the membrane attacking domain and β-hairpin P2 domains with respect to one another, allowing membrane insertion to take place^34^. Similar to these other toxins, a decrease in pH facilitated the MakA structural change comparable to the opening of a Swiss army knife blade when shifting from a soluble to a membrane-bound active state interaction with the lipid membrane. The pH-induced structural change potentiated the MakA-induced cytotoxic effect on the target cell under the conditions tested. Consistent with these results, we recently found that MakA co-localizes with β-catenin, actin and the phosphatidic acid biosensor, PASS, in filipodia-rich structures^21,22^.

Our *in vitro* analysis of the low-pH-induced interaction between MakA and host membranes included: purified lysosomes, cultured epithelial cells, red blood cells, and liposome models. In all model systems, we observed that low pH enhanced the activity of MakA, characterized by a striking tubulation of both purified lysosomes and liposomes prepared from epithelial cell total lipid extract. Furthermore, it was found that MakA formed tubular structures on liposomes independent of any specific protein receptor or energy-generating molecules, i.e., ATP or GTP, suggesting that at low pH the structural change and insertion of MakA itself was sufficient to trigger tubulation at the membrane surface. In contrast to some other PFTs that need cholesterol for oligomerization^35–37^, MakA could induce tubulation on liposomes obtained from lipid sources essentially lacking cholesterol, *i.e. E. coli* and *C. elegans*, showing that the pH-dependent tubulation of MakA can occur in the absence of cholesterol in the membrane. Bacterial cell membranes are typically devoid of cholesterol^38^, while the membranes of *C. elegans* are mostly composed of glycerophospholipids and sphingolipids, with trace quantities of cholesterol^39^. In addition, we observed that the presence of cholesterol was not required for the oligomerization of MakA in solution (**Supplementary Fig. 3b**).

Importantly, unlike Shiga toxin and cholera toxin, which insert into the membrane and cause inward-directed tubulations of artificial lipid membranes since toxin binding induces negative curvature of the plasma membrane^40–42^, MakA evidently could drive the rapid growth of tubules towards the extracellular space as shown with red blood cells (**Fig. 2g**). We propose that the appearance of tubular structures in response to MakA was a direct consequence of MakA insertion into the membrane, which appeared to create the conditions for generating a positive curvature, as described previously for protein-lipid complexes and multi-anchoring polymers^43,44^.

Our findings allowed us to propose a model for MakA oligomer assembly and its implications for pore formation. Within the oligomerized MakA filament, the TMHs line the inner cavity, interacting with lipids, which are intercalated both within the dimer interface and between the dimers in the helical spiral (**Fig. 4c** **and** **Fig. 5b**). In the context of pore-forming toxin systems, the initial insertion of one toxin component was described for XaxAB^17^, YaxAB^16^, and suggested for AhlC^19^. Considering that the TMHs of the MakA-tetramers harbor a lipid bilayer, it could be assumed that MakA initiates membrane insertion in a manner similar to its role in pore formation under neutral pH conditions when part of the tripartite MakA/B/E complex^13^. We hypothesize that under low-pH conditions, MakA transitions from inactive to the active stretched conformation and penetrates the membrane as a monomer. In the membrane bound state, the interaction of the head and tail domains of two monomers might lead to MakA dimer formation and subsequent oligomerization.

The pronounced structural difference between two subunits forming a dimer, in the absence of the tripartite complex interaction partner, i.e., MakB or MakE, would reflect how MakA’s plasticity nevertheless can structurally mimic the expected structure when involved in the tripartite cytolysin. First, one completely stretched monomer dimerizes with a monomer with a pronounced elbow-like kink from α3 to α4 in the transition from the tail to the neck region (**Fig. 5c**). Subsequently, the two dimers form a tetramer with the stretched and kinked MakA states associating with each other, respectively. The subsequent tetramerization followed by oligomerization does not result in pore formation, but helical growth and shearing off of lipids from vesicles and cell membranes at low pH. This process, which does not occur in the tripartite cytotoxin scenario under neutral pH conditions, depletes membranous structures of lipids and potentially causes cell lysis in a manner quite different from that of a *bona fide* α-PFT cytolysin complex. Our findings with the *V. cholerae* MakA protein reveal an unexpected capability and remarkable mode of action of an individual α-PFT toxin subunit.

## Materials and Methods

### Chemical and Lipids

Chloroform, formaldehyde, methanol, sodium citrate, Tween 20, Triton X-100 and Fluoromount were from Sigma (St Louis, MO, USA). Hoechst 33342 and Lysotracker were from Thermo Fisher Scientific (Waltham, MA, USA). Propidium iodide was from BD Biosciences (San Jose, CA). Protease inhibitor and phosphatase inhibitor, phosSTOP were from Roche (Roche AB, Solna, Sweden). All lipids were purchased from Avanti Polar Lipids (Alabaster, AL, USA). Lipids: 1-palmitoyl-2-oleoyl-glycero-3-phosphocholine (POPS), 1,2-dioleoyl-sn-glycero-3-phosphoethanolamine (DOPE), 1-palmitoyl-2-oleoyl-sn-glycero-3-phospho-L-serine (POPS), Sphingomyelin from Porcine Brain (SM), Cholesterol from Ovine (Chol), L-α-phosphatidylinositol-4,5-bisphosphate from Porcine Brain (PIP2), and N-palmitoyl-sphingosine-1-{succinyl[methoxy(polyethylene glycol)5000]} (PEG5Kce). Lyophilized PIP2 lipids were dissolved in a mixture of chloroform: methanol (2:1) to a concentration of 1 mg/mL. Next, they were protonated by addition of 0.5 μL of 1 M HCl to 100 μg of PIP2, kept at room temperature for 15 min and dried with nitrogen gas. The dried lipid was redissolved in chloroform: methanol (3:1) mixture to 1 mg/mL followed by drying again. Finally, the 100 μg of PIP2 was redissolved in 100% chloroform to 1 mg/mL and stored at –20°C until used for liposome production.

### Mammalian cell culture

Caco-2 (ATCC), HCT8 (ATCC) and HCT116 (ATCC) cells were cultured in RPMI-1640 medium (Sigma-Aldrich) supplemented with 10% fetal bovine serum (FBS), 1% penicillin/streptomycin, and non-essential amino acids. Cells were cultured at 37 °C, 5% CO_2_ and 90% humidity in 96-well plates overnight (for MTS cell viability assays), Coverslip bottom 8-well chamber slide (for confocal and spinning disc confocal microscopy) for 75-mm flasks (for lysosomes isolation), 24-well plates (for flow cytometry) and six-well plates (for Western blot analysis).

### Antibodies

Anti-MakA antiserum produced by GeneCust (1:20,000 dilution), anti-LAMP1 antibody (#9091) purchased from Cell Signaling (1:1000 dilution), and anti-beta-actin antibody (A2228) purchased from Sigma-Aldrich (1:5000 dilution) were used in this study.

### Cloning and purification of MakA

Cloning, overexpression, and purification of MakA have been previously reported^11^. AlexaFluor568 labeling of MakA was performed using an AlexaFluor568 protein labeling kit (Thermo Fisher) according to the manufacturer’s instructions.

### Isolation and treatment of intact epithelial cell lysosomes

HCT8 cells were grown overnight in RPMI-1640 complete media (∼pH 7.2). The following day cells were treated with MakA (250 nM, 18 h). At the end of treatment, lysosomes were purified from vehicle or MakA-treated HCT8 cells using Lysosome Isolation Kit (ab234047, Abcam), according to the manufacturer’s instructions.

For the lysosome pull-down assay, intact lysosomes freshly isolated from HCT8 cells were diluted in three times their volume of freshly prepared binding buffer (120 mM sodium citrate, pH 5.0, pH 6.5 or pH 7.0), followed by incubation with MakA (20 µg/mL) at 37°C (60 min). These MakA-lysosome complexes were centrifuged at 21,000 x g (30 min, RT), pellets were washed in the respective binding buffer and resuspended in 2X Laemmli buffer. Samples were run on an SDS-PAGE, and after electrophoresis, the samples were transferred to a nitrocellulose membrane. A Western blot analysis was performed using anti-MakA antiserum (1:20,000 dilution, overnight at 4°C) that was detected with HRP-conjugated goat anti-rabbit secondary antibodies. Detection of LAMP1, using anti-LAMP1 antibodies, was used as an internal loading control for lysosome pull-down experiments. The membranes were developed with a chemiluminescence reagent (Bio-Rad). Images were acquired using an ImageQuant™ LAS 4000 instrument and processed using ImageJ-FIJI distribution^45^.

For confocal microscopy, intact lysosomes diluted in three times their volume of freshly prepared binding buffer (120 mM sodium citrate, pH 5.0, pH 6.5 or pH 7.0) were exposed to Alexa568-MakA (1 µM). To facilitate the binding, Alexa568-MakA and lysosomes were incubated in a 37°C incubator for 60 min. At the end of treatment, samples were visualized by a Leica SP8 inverted confocal system (Leica Microsystems) equipped with an HC PL APO 63x/1.40 oil immersion lens. Images were captured using LasX software (Leica Microsystems) and processed using ImageJ-FIJI distribution^45^.

### Live cell spinning disk microscopy

Live cell experiments were conducted in phenol-red-free IMDM media adjusted to pH 5.0 supplemented with 10% FBS and 1 mM sodium pyruvate (Thermo Fisher Scientific) at 37°C in 5% CO_2_. Alexa568-MakA (500 nM) was added to HCT8 cells, and images were recorded every 1 min during a period of 120 min using a 63X lens and Zeiss Spinning Disk Confocal controlled by ZEN interface (RRID:<u>SCR_013672</u>) with an Axio Observer Z1 inverted microscope, equipped with a CSU-X1A 5000 Spinning Disk Unit and an EMCCD camera iXon Ultra from ANDOR. Images were processed with Zeiss ZEN Lite and ImageJ-FIJI distribution^45^.

### Immunofluorescence

Lysosomal tubulation was investigated by treating Caco-2 cells with Alexa568-MakA (250 nM, 18 h) in IMDM complete media (pH 7.2). After treatment, cells were subsequently counterstained for lysotracker (200 nM, 30 min) and Hoechst 33342 (2 μM, 30 min).

For confocal microscopy, HCT8 cells were loaded with the nuclear staining marker Hoechst 33342 (2 μM, 30 min) and exposed to Alexa568-MakA (500 nM) in different pH-adjusted IMDM complete media for 4 h at 37°C in 5% CO_2_. Cells were visualized live using a Leica SP8 inverted confocal system (Leica Microsystems) equipped with an HC PL APO 63x/1.40 oil immersion lens. Images were captured using the LasX (Leica Microsystems) and processed using ImageJ – FIJI distribution^45^.

For the propidium iodide uptake experiment, HCT8 cells were treated with MakA (500 nM, 4 h) in IMDM complete media (pH 5.0 or pH 7.4), followed by adding propidium iodide (0.5 μg/mL, 30 min). Fluorescence and bright-field images were captured with a fluorescence microscope (Nikon, Eclipse Ti). Images were processed using the NIS-Elements (Nikon) and ImageJ – FIJI distribution^45^.

For Alexa568-MakA binding to erythrocytes, freshly prepared human erythrocytes (0.25% in PBS) were loaded into an 8-well chamber slide (µ-Slide, ibidi), cells were allowed to adhere to the glass surface for 10 h, followed by buffer exchange to citrate buffer (pH 5.0, pH 6.5 or pH 7.4). The erythrocytes were exposed to Alexa568-MakA (500 nM) for 3 h at 37°C in 5% CO_2_. Cells were visualized using a Leica SP8 inverted confocal system (Leica Microsystems) equipped with an HC PL APO 63x/1.40 oil immersion lens. The maximum z-stack projection of the human erythrocytes treated with Alexa568-MakA (pH 6.5 in citrate buffer) was constructed using Leica LasX Software.

### Cell toxicity assay

HCT8, Caco-2 and HCT116 cells were treated with increasing concentrations of MakA at a given pH at the indicated time point (4 h or 24 h). At the end of treatment, cells were incubated with MTS reagent (15 min) at 37°C in an incubator, and cell viability was quantified by measuring MTS absorbance on an Infinite M200 microplate reader (Tecan). Data were normalized to the vehicle-treated cells (pH 7.2) and expressed as a percentage of the control.

For flow cytometry experiments, HCT8 cells were grown on a 24-well plate (8×10^4^/well, Tecan Group Ltd) overnight in IMDM complete media (pH 7.2). The following day, cells were treated with vehicle (Tris 20 mM) or MakA (500 nM, 4 h) in media adjusted to a given pH. At the end of treatment, cells were incubated with propidium iodide (0.5 μg/mL) at 37°C for 30 min. Cellular uptake of propidium iodide in vehicle or MakA-treated cells was investigated by flow cytometry. Cellular uptake of propidium iodide was quantified and presented as mean fluorescence intensity (MFI) for the gated cells.

### Human erythrocytes hemolysis assay

Freshly prepared human erythrocytes (0.25%) in citrate buffer (120 mM sodium citrate, pH adjusted to 5.0, 6.5 or 7.4) were added to a 96-well plate. The erythrocytes were treated with increasing concentrations of MakA at two different time points (90 min and 5 h) at 37°C in 5% CO_2_. After centrifugation (500 x g), the supernatants were monitored spectrophotometrically for released hemoglobin by measuring absorbance at 545 nm to indicate red blood cell lysis. MakA-induced hemolysis of erythrocytes was normalized against erythrocytes treated with Triton X-100 (0.1%). Data were expressed as a percentage.

### Scanning Electron Microscopy

Freshly prepared human erythrocytes (0.25%) were treated with MakA (500 nM, 90 min) in citrate buffer (120 mM, pH 6.5). Samples were fixed with a fixative (1% glutaraldehyde + 0.1 M CaCo buffer + 3 mM MgCl_2_) in the microwave and washed twice with buffer (0.1 M CaCo buffer + 2.5% sucrose + 3 mM MgCl_2_). They were sedimented onto poly-L-lysine-coated coverslips for 1 h and subsequently dehydrated in series of graded ethanol solutions in the microwave. The samples were then dried to a critical point (Leica EM300). Subsequently, samples were coated with 2 nm of platinum (Quorum Q150T ES). Samples were imaged with field-emission scanning electron microscopy (FESEM; Carl Zeiss Merlin) using an in-chamber (ETD) secondary electron detector at an accelerating voltage of 5 kV and probe current of 150 pA.

### Extraction of epithelial cell lipids for liposome binding assays

Lipids were extracted by the Folch method^46^ from 10 x 150 cm^2^ confluent flasks of HCT8 cells. Briefly, the HCT8 cell lipid extracts, dissolved in chloroform, were dried to a thin film under a nitrogen stream. The dried lipid yield was 12 mg. The lipid film (5 or 10 mg/mL) was hydrated in HEPES buffer (10 mM HEPES, 150 mM NaCl, pH 7.4), citrate buffer (120 mM citrate buffer, pH 6.5) or (20 mM citric acid, 50 mM KCl, 0.1 mM EDTA, pH 4.5). The lipid suspension was extruded through polycarbonate membranes (0.1 μm) using the Avanti Mini-Extruder (Avanti Polar Lipids, Alabaster, AL, USA).

The liposome pull-down assay was performed as previously described^21,47^. Briefly, the liposome suspension was diluted in five times its volume of freshly prepared binding buffer (120 mM sodium citrate, pH 5.0, pH 6.5 or pH 7.4), followed by centrifugation at 21,000 x g for 30 min at room temperature. The liposome pellet was resuspended in binding buffer followed by incubation with MakA (20 µg/mL). The liposome-protein mixtures were incubated at 37°C (60 min), followed by centrifugation at 21,000 x g at room temperature (30 min). To reduce the background, pellets were washed in the respective binding buffer two to three times. The resulting sample was loaded onto the SDS-PAGE, transferred to a nitrocellulose membrane and subjected to Western blot analysis using anti-MakA antiserum (1:10,000 dilution, overnight at 4°C). The MakA antibodies were detected with HRP-conjugated goat anti-rabbit secondary antibodies. The membranes were developed with a chemiluminescence reagent (Bio-Rad). Images were acquired using ImageQuant LAS 4000 instrument and processed using ImageJ - FIJI distribution^45^.

### Circular dichroism spectroscopy

Far-UV Circular Dichroism (CD) analysis of MakA protein or MakA and ECLE liposome complexes was performed using Jasco J-720 Spectropolarimeter (Japan) at 25°C. Briefly, MakA (3 µM) alone or MakA (3 µM) and ECLE (5 mg/mL) were incubated overnight at 25°C in citrate buffer (5 mM) with varying pH. The spectra were recorded between 195-260 nm using 2-s response time, a 1 mm cuvette path length and a 2 nm bandwidth. Data of an average of five repeated scans were used for graphical presentation and analyses.

### Liposome preparation

Liposomes containing 0.5 mol% PEG5Kce, 5 mol% PIP2, 10 mol% SM, 10 mol% POPS, 15 mol% DOPE, 20 mol% Chol, and 39.5 mol% POPC (referred to herein as synthetic lipid mixture [SLM] liposomes) were prepared using the lipid film hydration and extrusion method. SLM+TxRed liposomes were created using the same protocol as above except for the addition of 1 mol% TxRed-DHPE and the corresponding reduction of POPC content to 38.5 mol%. The individual lipids dissolved in chloroform were mixed together, dried under nitrogen and then stored in a vacuum for a minimum of one hour. The dried lipid film was then rehydrated using a pre-heated citrate-potassium buffer at pH 4.5 (20 mM citric acid, 50 mM KCl, pH 4.5, 40°C) to a lipid concentration of 1 mg/mL. The solution was then extruded at ∼40°C eleven times through a polycarbonate membrane with 100-nm pore size using an Avanti mini extruder. The liposomes were stored at 4°C until used.

### Fluorescence microscopy of SLBs

Supported lipid bilayers (SLBs) were formed on glass coverslips. Coverslips were cleaned by boiling in 7X detergent (MP biochemicals) for 2 h followed by extensive rinsing in 18 MΩ water and blown dried with nitrogen. Clean coverslips were fitted with Poly(dimethylsiloxane) (PDMS) sheets containing 10 µm holes to create glass-bottomed wells. SLBs were made by adding 10 µL of 0.1 mg/mL of either SLM or SLM+TxRed liposomes to each well and incubating the wells at 37°C for 30 min before rinsing the wells with citrate-potassium buffer to remove excess liposomes. Wells were then extensively rinsed with citrate buffer at pH 6.5 (120 mM sodium citrate) prior to protein addition. Either Alexa568-MakA or unlabeled MakA diluted in citrate buffer at pH 6.5 was then added to a well to reach a final concentration of 3 µM. The SLB surface was monitored using a Nikon Eclipse Ti2-E inverted epifluorescence microscope equipped with a 60X objective multi-band pass filter cube 86012v2 DAPI/FITC/TxRed/Cy5 (Nikon Corp.), Prime 95B sCMOS camera (Teledyne Photometrics), and Spectra III light source (Lumencor).

### Surface plasmon resonance

To analyze the interaction of MakA with ECLE or POPC liposomes, L1 sensor chips and a Biacore 3000 instrument were used as previously described^21^. In brief, the ECLE or POPC liposomes were immobilized onto the cells of the L1 sensor chip surfaces at low flow rates of 2 μL/min for 40 min, stabilized with 50 mM NaOH, and the successful surface coverage was tested by injecting Bovine Serum Albumin (BSA). After successful surface coverage, two-fold serially diluted MakA (diluted in the SPR running buffer (120 mM citrate buffer, pH 6.5) with increasing concentrations (0 to 200 nM) was injected for 120 sec at flow rates of 5 μL/min. For the binding analysis with POPC liposomes, the maximum concentration of MakA (200 nM) was used. All experiments were repeated at least twice, and the backgrounds of control flow cells were subtracted from the experimental cells before final data processing. The binding affinities (KD) were determined from the concentration gradient experiments, and the binding responses at equilibrium were fit to a simple 1:1 steady-state affinity model using the global data analysis option available within the Scrubber 2 software (http://www.biologic.com.au/scrubber.html).

### Western blot analysis

For Western blot analysis, HCT8 cells were grown on a 6-well slide (3×10^5^/well, Thermo Scientific) overnight. The pH of IMDM cell culture media supplemented with 10% FBS and 1% penicillin/streptomycin was adjusted to either 5.0, 6.5, 7.4 or 8.0 followed by treatment with an increasing concentration of MakA for 4 h. Cells were rinsed with ice-cold PBS to remove unbound MakA and lysed in ice-cold NP-40 cell lysis buffer (20 mM Tris-HCl pH 8, 0.25% Nonidet P-40, 10% glycerol, 0.5 mM EDTA, 300 mM KCl, 0.5 mM EGTA, 1x phosSTOP, and protease inhibitor cocktail from Roche). After mixing with sample buffer, cell lysates were boiled for 5 min and separated by SDS-PAGE. The proteins were transferred to a nitrocellulose membrane and blocked with 5% skim milk in 0.1% PBST (RT, 1 h). The membranes were incubated with respective primary antibodies in 5% skim milk (4°C, overnight). After washing with PBST (0.1%), membranes were incubated with HRP-conjugated secondary antibodies in 5% skim milk (RT, 1 h). The membranes were developed with Immun-Star™ AP Chemiluminescence (Bio-Rad). Images were acquired using ImageQuant LAS 4000 instrument and processed using ImageJ - FIJI distribution^45^.

### Transmission electron microscopy

Negative staining for lysosomes or liposomes was performed on glow discharged copper grids (300 mesh) coated with a thin carbon film (Ted Pella, Redding, CA). After adding 3 μL sample to the grids, they were washed twice with MQ water and stained with 1.5% uranyl acetate solution (EMS [Hatfield, PA]), followed by MQ water washing. Grids were examined with Talos L120C, operating at 120 kV. Transmission electron micrographs (TEM) were acquired with a Ceta 16M CCD camera using TEM Image & Analysis software ver. 4.17 (FEI, Eindhoven, The Netherlands).

### Cryo-electron microscopy sample preparation and data collection

The ECLE liposomes (10 mg/mL) were incubated with MakA (30 μM) in binding buffer (120 mM sodium citrate, pH 6.5) for 60 min at 37°C. Quantifoil 2/1-200 grids were glow discharged before the addition of 3 μL protein-liposome mixture. Grids were then flash-frozen in liquid ethane using an FEI Vitrobot (Thermo Fisher Scientific). Data collection was performed at the Umeå University Core Facility for Electron Microscopy (UCEM) on a Titan Krios (Thermo Fisher Scientific), operating at 300 kV and equipped with a Gatan K2 BioQuantum direct electron detector (Gatan, Inc.). Images were acquired using EPU (Thermo Fisher Scientific). A total of 2,476 movies, each with 40 frames over a total dose of 43 e-/Å^2^, and a 0.75 to 2.5 μm defocus range at a 1.042 Å pixel size were collected.

### Cryo-EM data processing and helical reconstruction

The MotionCorr implementation of RELION-3.1 was used for drift correction and dose weighting of the micrographs^48^. The contrast transfer function (CTF) was determined using CTFFIND-4.1.14 (ref.^49^), and empty or micrographs with poor CTF fits or low ice quality were removed after manual inspection, which reduced the total to 1,351 micrographs (**Supplementary Fig. 4**). Helical reconstruction tools in RELION-3.1 (ref. ^50^) were used for subsequent image processing. Filaments were selected manually using the helix picker in RELION-3.1 with an outcome of 13,784 picked start and endpoints. First, segments were extracted between the picked start and endpoints as 2x binned data using a box size of 437 Å (210 pixels, 2.084 Å/px) with an inter-box distance (IBD) of 23 Å, resulting in 195’809 segments, which were subjected to 2D classification with 80 classes and a spherical mask of 360 Å. Classes displaying a straight filament with high-resolution features (152’961 segments) were refined without symmetry using a featureless cylinder (diameter 320 Å) generated with the helix toolbox in RELION-3.1 (**Supplementary Fig. 4**). In parallel, a 2D classification was performed by extracting 65’241 segments from 2x binned data, applying a box size of 646 Å (310 px, 2.084 Å/px), an IBD of 62 Å, and a spherical mask of 580 Å. The volume refined without symmetry and the 2D class averages obtained from large helical segments were used to determine the helical symmetry parameters. Diameter and repeat distance were visually analyzed and measured in a representative 2D class average in RELION-3.1 (**Fig. 4b** **and Supplementary Fig. 5a**). Additionally, the repeat distance was calculated from the corresponding collapsed power spectrum (layer-line distance-1) in SPRING-0.68 (ref. ^51^) (**Supplementary Fig. 5b**). Next, to determine the helical twist and rise, the number of turns and subunits per repeat were counted from the initial reconstructed model (**Supplementary Fig. 5c-d**), and the handedness of the reconstruction was, after the subsequent high-resolution refinement described below, confirmed using the MakA crystal structure^11^.

For the final reconstruction, 95’603 segments were extracted unbinned with 460-pixel boxes (479 Å) and an IBD of 46.56 Å. Subsequent 2D classification with a 432 Å spherical mask resulted in 65’485 segments that were 3D refined using a featureless cylinder as a template and a spherical mask of 360 Å. Local symmetry searches were performed to narrow down the helical symmetry by refining helical twist (48-50°) and rise (5.4-6 Å), yielding a map with an overall 4.1 Å resolution. Subsequent Bayesian polishing improved the resolution to 3.8 Å, and estimation of anisotropic magnification and CTF refinement resulted in a final map with an overall resolution of 3.7 Å (**Supplementary Fig. 6**). As the peripheral region was less well-resolved, a subsection of the structure was isolated via signal subtraction, centered in a 260-pixel box and subjected to 3D classification without symmetry and local searches with increasing sampling rate from 3.7°, 1.8°, and 0.9° angles. From the resulting three classes, further refinement of class 2 (56.8%) yielded a map of the two tetramers in isolation at an overall resolution of 4.1 Å with improved peripheral density (**Supplementary Fig. 4**, blue branch, **and Supplementary Fig. 6d,e**). As the resulting three classes showed different conformations of the MakA head region, we examined whether these conformations exist across the volume. Two subsequent 3D classifications into five classes, without image alignment but with local symmetry searches, first with all classes, then with the top class from the first run, showed a normal distribution of angles, ranging from 48.48° to 48.68° suggesting continuous motion/rotation along the filament axis, which is most pronounced in the tail region (**Supplementary Fig. 4,** blue branch, lower right).

### Model building, refinement, and validation

To obtain an initial model of the tail domain, the MakA crystal structure (PDB-6EZV^11^) was rigid-body docked into the density map using Chimera^52^ and Coot^53^. Regions where the density/model fit was poor (no density, difference in conformation) were trimmed. This included the C-terminal tail (res. 351-365) and the central region (res. ∼160-260). Elements in the tail domain with poor density fit were rigid-body docked. The central region of the protein, which includes the neck and head, was built *de novo* in Coot^53^. The model was first refined against the asymmetric map of the two tetramers in isolation (4.1 Å) using phenix.real_space_refine (version 1.14-3260)^54^. Next, this model was rigid-body docked into the 3.7-Å helical map, two neighboring placeholder molecules were symmetry expanded to provide interaction interfaces, and refined with secondary structure restraints. The final model contains four MakA subunits with trimmed sidechains in the tail domain (N-terminus to 159, 281 to C-terminus, **Supplementary Fig. 6e,f**). To validate the final model, all atomic coordinates were displaced randomly by 0.5 Å, refined against half map 1, followed by calculating the Fourier-Shell Correlation coefficient of the resulting refined model, and half map 1 or half map 2 (ref ^55^). Model statistics are presented in (**Supplementary Table S1**).

### Map and model visualization

Structure analyses and preparation of the figures were performed using PyMOL (Schrödinger) or UCSF ChimeraX^56^.

### Statistical analysis

The result from replicates is presented as mean ± s.e.m. or mean ± s.d. The statistical significance of different groups was determined by Student’s t-tests (two-tailed, unpaired) or one-way ANOVA using Microsoft Excel or GraphPad Prism. *p ≤ 0.05, **p ≤ 0.01, ns = not significant.

## DATA AVAILABILITY

The cryo-EM density maps have been deposited in the EM Data Bank with accession code EMD-13185 (MakA helical reconstruction) and EMD-13185-additional map 1 (two tetramers refined in isolation). Coordinates have been deposited in the Protein Data Bank under accession code PDB-7P3R.

## ACKNOWLEDGMENTS

This work was supported by grants from the Swedish Research Council (No. 2018-02914 to S.N.W.; No. 2016-05009 to K.P; No. 2019-01720 to B.E.U.; No. 2016-06963 to G.G.), The Swedish Cancer Society (No. 2017-419 and No. 2020-711 to S.N.W.), The Kempe Foundations (No. JCK-1728 to S.N.W.; No. SMK-1756.2 and No. SMK-1553 to K.P.; No. JCK-1724 and No. SMK-1961 to B.E.U), and the Faculty of Medicine, Umeå University (Strategic Research Grant 2019-2021 to S.N.W.). M.B. was supported by the Knut and Alice Wallenberg Foundation. J.B. acknowledges funding from the Swedish Research Council (2019-02011), the European Research Council (ERC Starting Grant PolTube 948655), the SciLifeLab National Fellows program, and MIMS. We acknowledge the Protein expression and purification facility (PEP) at Umeå University for construct design and cloning. We acknowledge the facilities and technical assistance of the Umeå Core Facility Electron Microscopy (UCEM) and the Biochemical Imaging Center (BICU), Umeå University, a part of the National Microscopy Infrastructure NMI (VR-RFI 201600968 and VR-RFI 2019-00217). The CryoEM data were collected by UCEM, which is a node of the Swedish National Cryo-EM Facility, funded by the Knut and Alice Wallenberg Foundation, Erling-Persson Family Foundation, The Kempe Foundations, SciLifeLab, Stockholm University, and Umeå University.

## AUTHOR CONTRIBUTIONS

A.N. performed confocal microscopy, transmission electron microscopy, cryo-electron microscopy sample preparation for data collection, flow cytometry and cell toxicity; K.P. purified protein; E.T. and S.L.M, assisted A.N. with Western blot. A.N., H.P., E.T., At. A. and M.B. performed liposome experiments. At. A., and J.Å., performed CD experiments. A.B. and J.B. performed cryo-EM data processing and analysis. N. Z. assisted A.N. with hemolytic assay and analyzed the *mak* operon. A.N. wrote the initial version of the manuscript. All authors read and commented on the manuscript. M.B., K.P., A.S., G.G., B.E.U., and S.N.W. obtained the funding. A. N., M.B., K.P., B.E.U, and S.N.W. supervised the research and finalized the manuscript.

---

**Supplementary Fig. 1:**
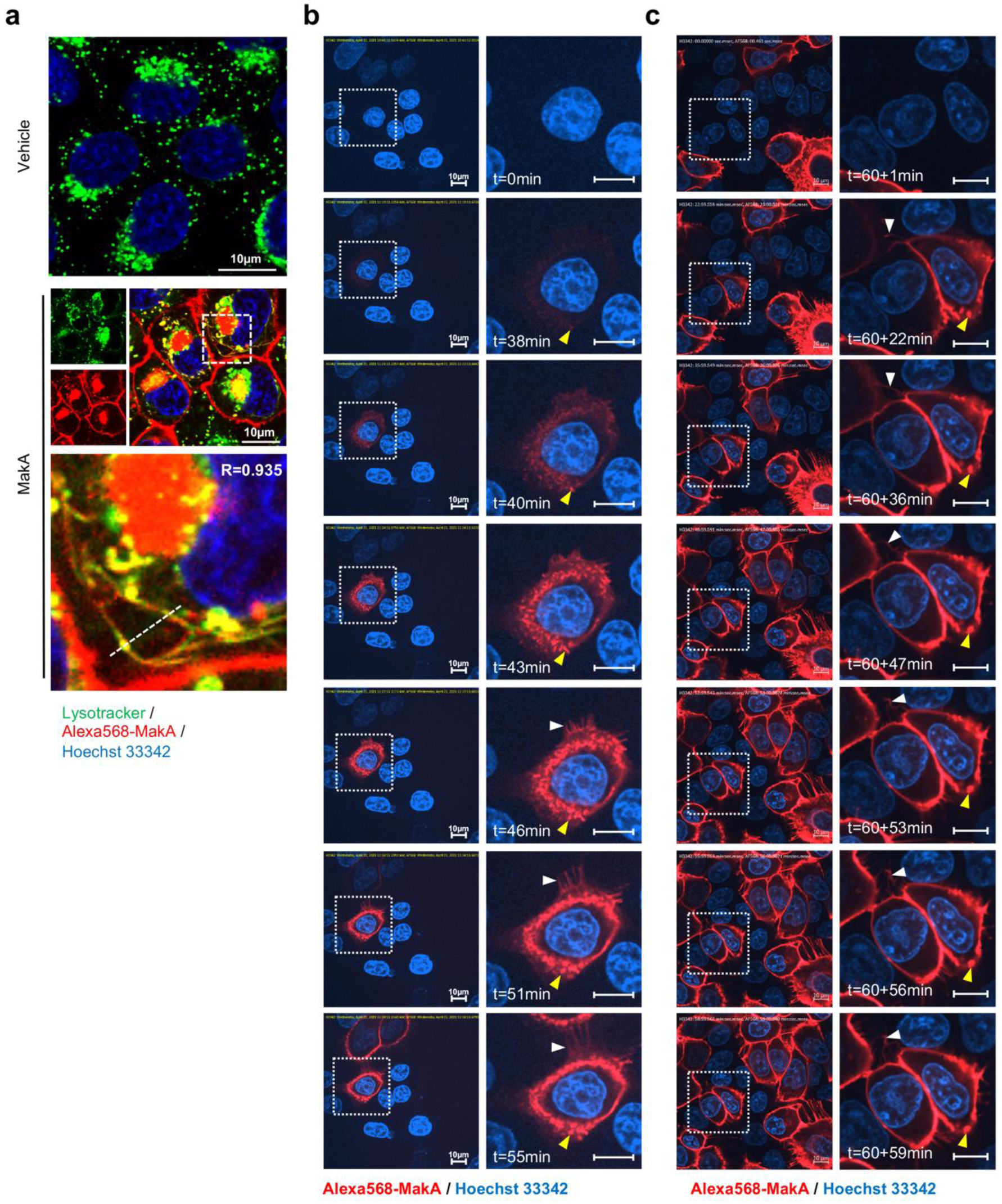
MakA binding to the epithelial cell membrane in filipodia rich structures. **a** Caco-2 cells treated with vehicle or Alexa568-MakA (250 nM, 18 h) and subsequently counterstained with Lysotracker (green, 500 nM, 30 min). Nuclei were counterstained with Hoechst 33342. Pearson correlation co-efficient was used for the calculation of Alexa568-MakA (red) co-localization with lysotracker (green) along the dotted line. Scale bars, 10 μm. **b-c** Still images of HCT8 cells exposed to Alexa568-MakA (500 nM) at pH 5.0. Yellow arrowheads indicate the accumulation of MakA in filipodia rich structures and white arrowheads indicate the appearance of MakA positive tubular structures. Nuclei were counterstained with Hoechst 33342. Scale bars, 10 μm. Images in (a) were acquired for a time limit of 55 min immediately after Alexa568-MakA administration, while images in (b) were acquired for 59 extra minutes, 60 min (Total time = 119 min) after Alexa568-MakA administration.

**Supplementary Fig. 2:**
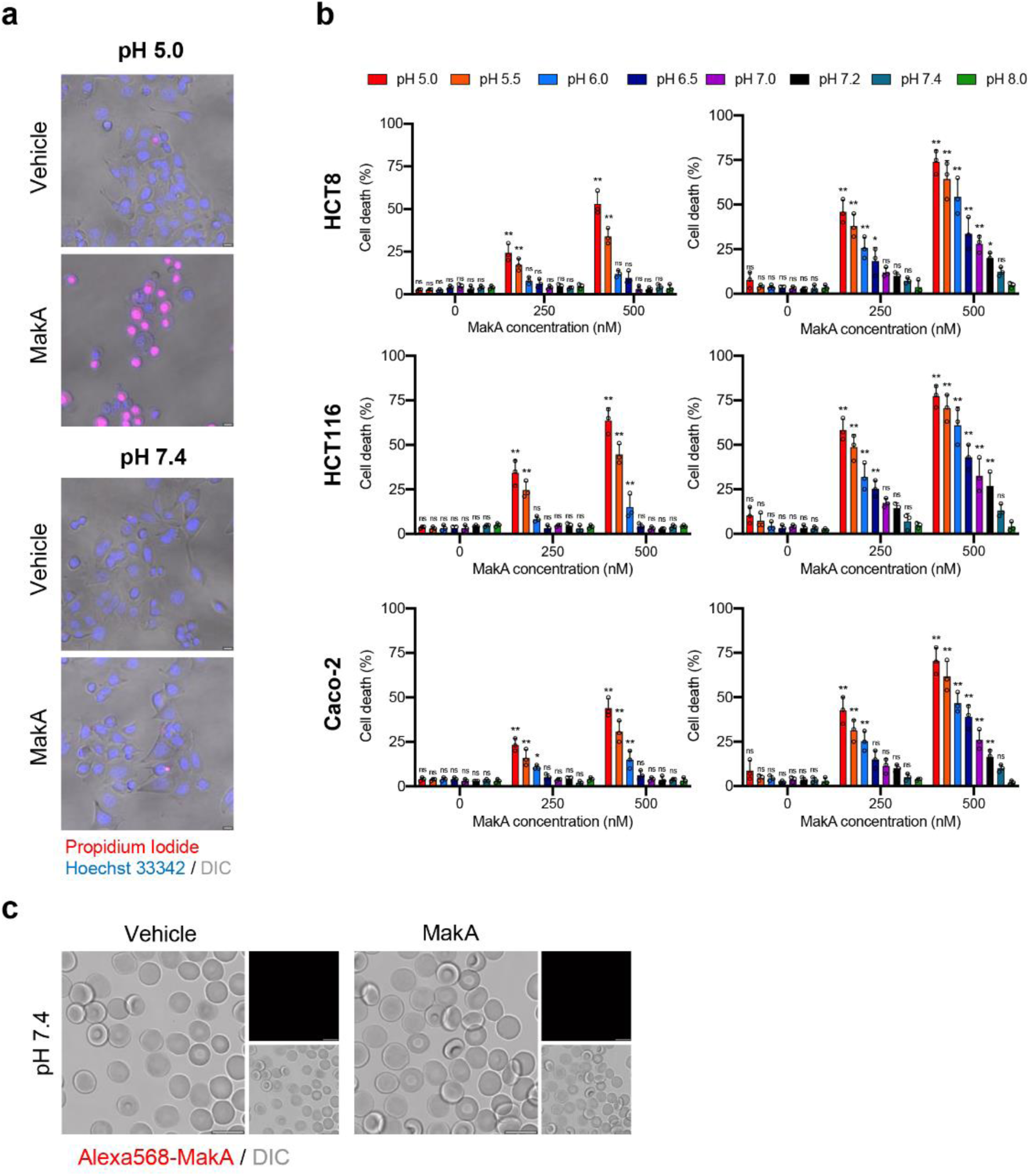
pH-dependent cytotoxicity of MakA in target cells. **a** MakA caused pH-dependent permeability of HCT8 cells as determined by cellular uptake of propidium iodide. The uptake of propidium iodide (red) was assessed by confocal microscopy. Scale bars, 10 µm. **b** MakA induced death of Caco-2, HCT116 and HCT8 cells under different pH conditions. Epithelial cell viability was assessed using the Trypan blue exclusion method. Data are representative of three independent experiments; bar graphs show mean ± s.d. Significance was determined from biological replicates using two-way analysis of variance (ANOVA) with Tukey’s multiple comparisons test. *p≤0.05, **p≤0.01, ns = not significant. **c** Human erythrocytes (0.25 %) in phosphate buffered saline (PBS) were allowed to adhere on the glass surface for 10 h, followed by buffer exchange to citrate buffer (pH 7.4). The erythrocytes were treated with Alexa568-MakA (500 nM, 3 h), and cell-bound MakA was detected by confocal microscopy. Scale bars, 10 µm.

**Supplementary Fig. 3:**
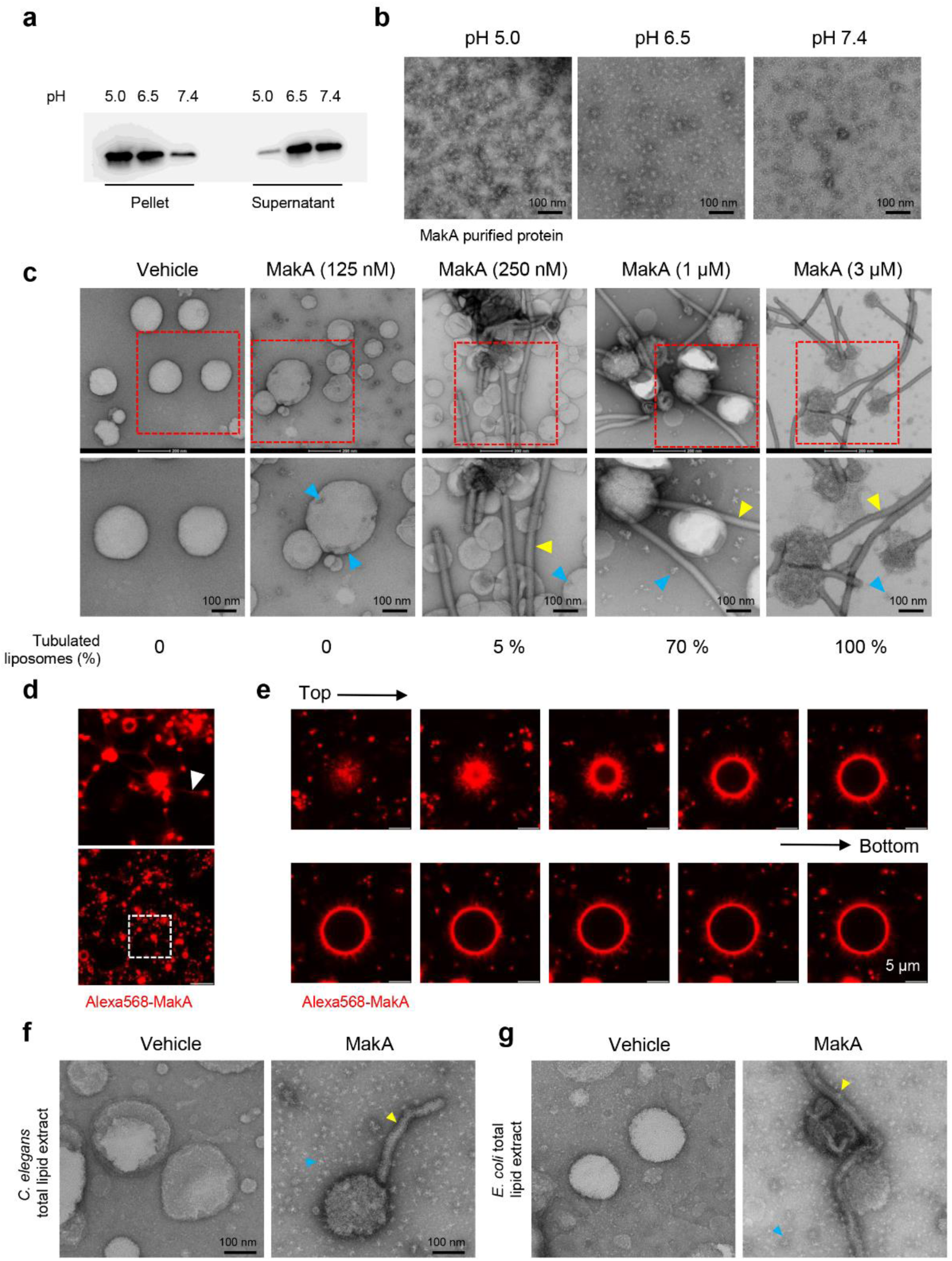
MakA binding to ECLE liposomes and induction of tubulation in a pH-dependent manner. **a** Western blot analysis of ECLE liposome-bound MakA detected with anti-MakA antisera. MakA protein (128 nM) was incubated under different pH conditions with liposomes prepared from epithelial cell lipid extract. S = supernatant and P = pellet. **b** EM micrographs of MakA protein (10 µM) that was incubated at 37°C for 1 h in 120 mM citrate buffer, adjusted to pH 5.0, pH 6.5 and pH 7.4, respectively. The protein samples were spotted on grids and stained with 1.5 % uranyl acetate solution. Images were captured with transmission electron microscopy (TEM). MakA appeared as oligomers of varying size under the different pH conditions. Scale bars, 100 nm. **c** ECLE liposomes were treated with increasing concentration of MakA for 90 min and stained with a 1.5 % uranyl acetate solution. Images were captured with TEM. Yellow arrowheads indicate tubular structures, blue arrowheads indicate the formation of MakA oligomeric structures present nearby or on liposomes. Scale bars, 200 nm and 100 nm. A quantification (%) of liposomes with tubular structures is shown below the micrographs. **d-e** ECLE liposomes were treated with Alexa568-MakA (1 µM, 90 min, pH 6.5). Liposome-bound MakA across the tubular structure (arrowhead, white) was detected by confocal microscopy. Scale bars, 5 µm. Selected images from z-stack projection of liposome-bound Alexa568-MakA are shown (Top = topmost section, Bottom = section close to the coverslip). The fraction of large vesicles was less than 1% in the reaction mixture. **f-g** Liposomes prepared from *C. elegans* or *E. coli* total lipid extracts were treated with MakA (3 µM, 90 min) at pH 6.5. Yellow arrowheads indicate MakA induced tubular structures and blue arrowheads indicate MakA oligomers. Scale bars, 200 nm and 100 nm.

**Supplementary Fig. 4:**
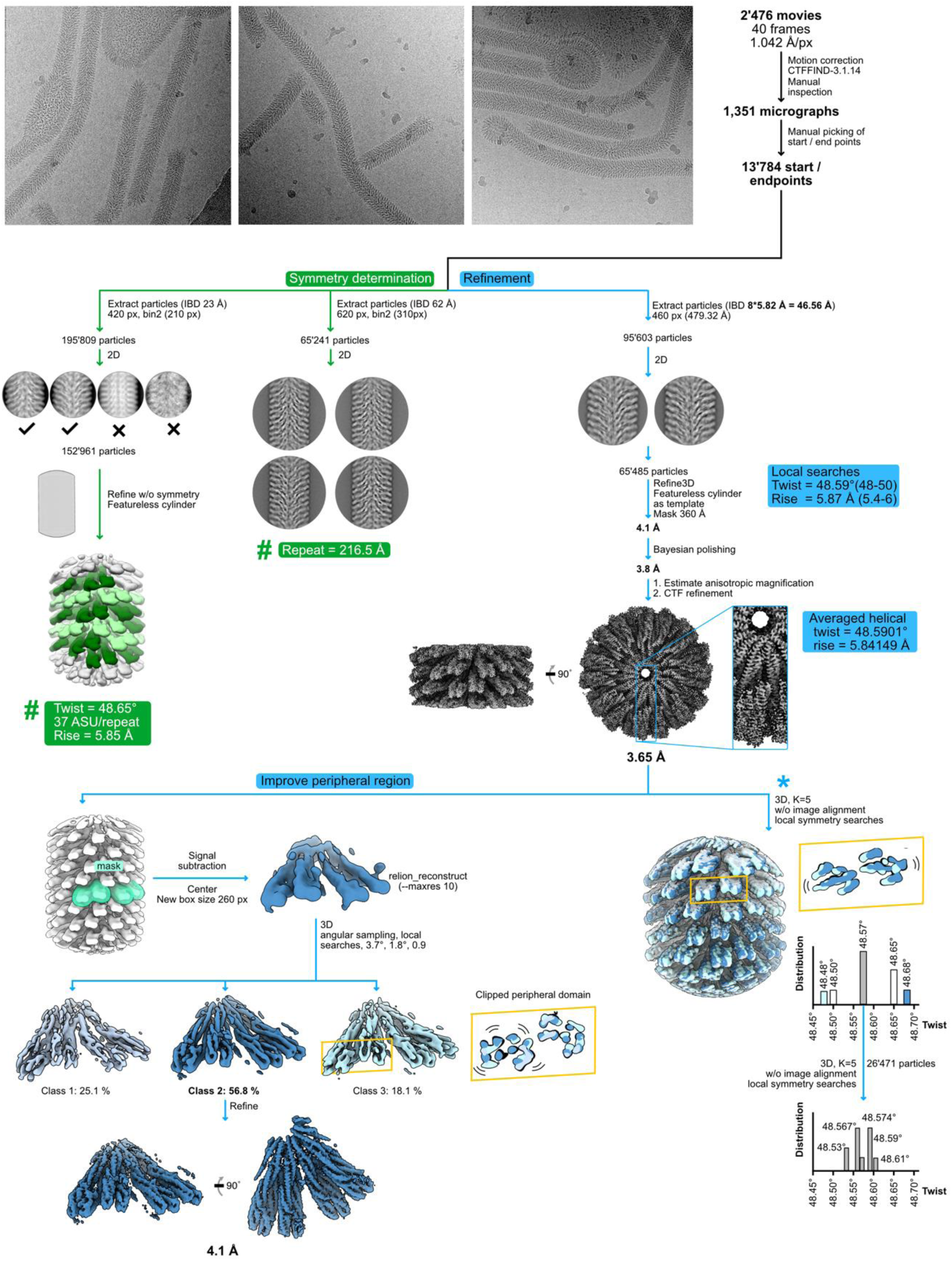
Cryo-EM processing scheme. Three representative micrographs are shown next to a schematic sorting and data processing tree. The green branch of the tree indicates steps performed to obtain symmetry parameters of the filament, while the blue branch outlines high-resolution refinement. Within the green branch, the (#) refers to (Supplementary Fig. 5), which details how the listed symmetry parameters were obtained. The branch depicted with a (*) describes two consecutive 3D classification experiments, performed without image alignment but with local searches of helical symmetry, to analyze the presence of continuous motion or concrete states in the filament.

**Supplementary Fig. 5:**
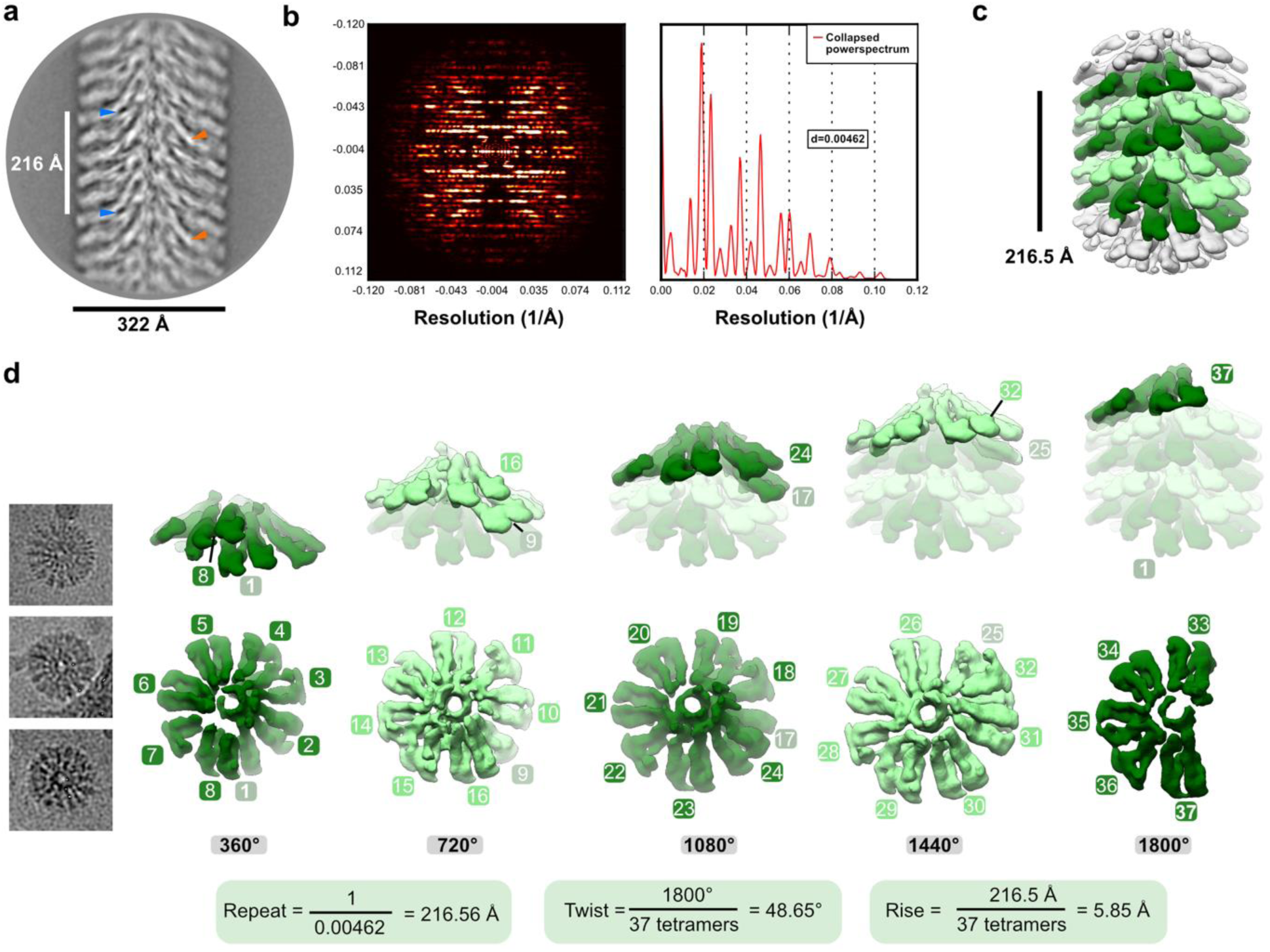
Helical symmetry determination of the MakA-filament. **a** A representative 2D class average, obtained from segments extracted with a large box size of 646 Å using RELION-3.1 (ref. ^1^), is shown. Visual analysis and measurements of the distance between repetitive elements in the 2D class averages (examples indicated with blue or orange arrows) suggest a repeat distance of 216 Å. **b** Enhanced (left), and corresponding collapsed power spectrum (right, red curve), obtained from the class average shown in (a) using SPRING-0.86 (ref. ^2^), confirms a repeat distance of ∼216 Å. **c** A volume, obtained by refining well-defined class averages showing high resolution features without symmetry in RELION-3.1, is shown 20 Å low-pass filtered with one repeat colored in shades of green. **d** Three filament top views, cut out from micrographs, are shown next to a dissection of the volume depicted in (c). The five turns, constituting one repeat, are demonstrated individually, starting with the first (left) and ending with the last turn (right). The lower row visualizes a top view of each turn with individual repetitive segments numbered. The top row shows the corresponding turns in a side view in solid colors, superimposed with a transparent filament. The resulting calculated symmetry parameters are shown below the dissected repeat.

**Supplementary Fig. 6:**
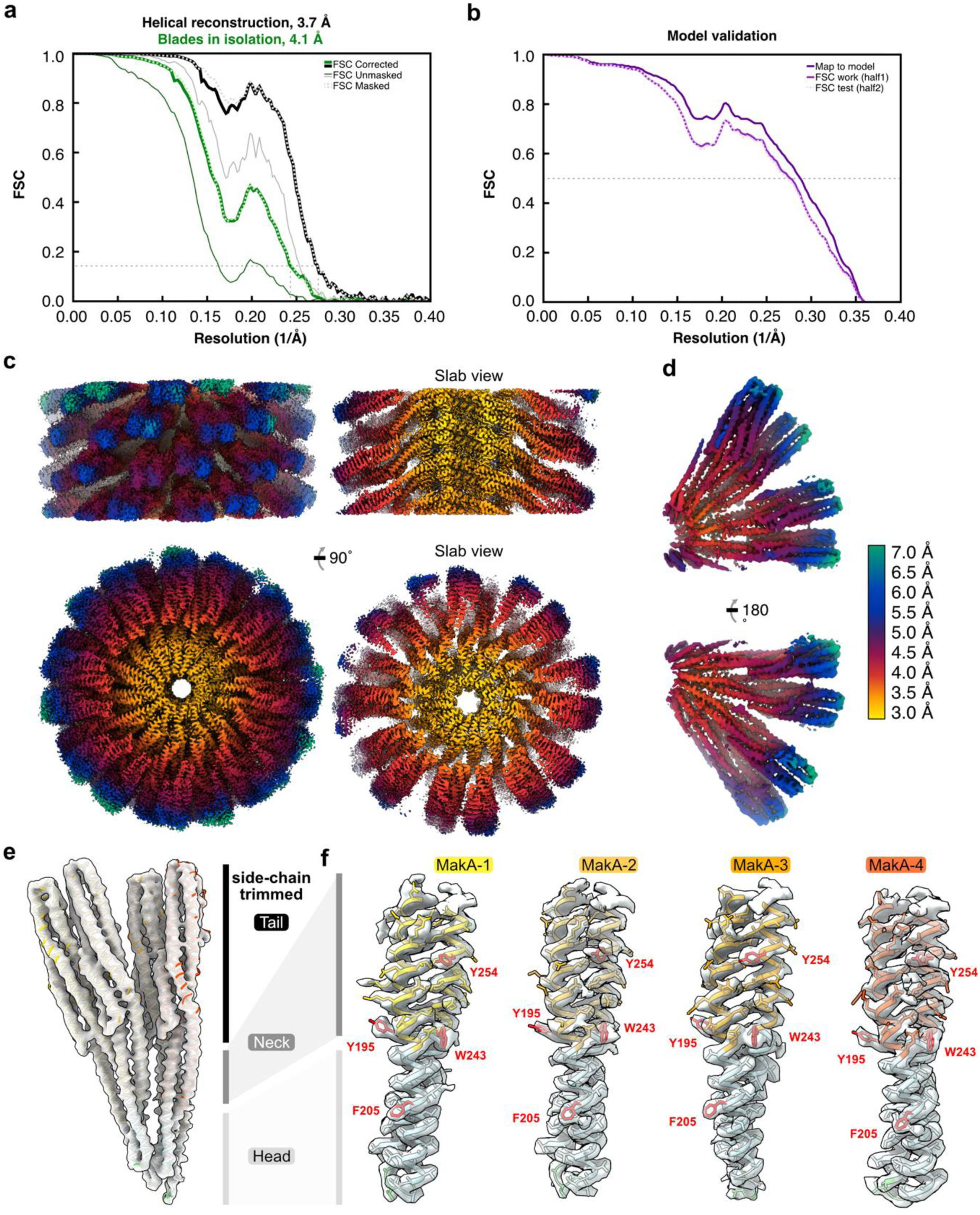
Global and local resolution estimation, model validation, and density fit. **a** The Fourier Shell Correlation (FSC) curves, obtained from RELION-3.1 (ref. ^1^), are shown for the helical reconstruction (EMD-13185) and the two tetramers refined in isolation (EMD-13185-additional map 1). An FSC value of 0.143 is indicated with a thin dashed line. **b** Cross-validation of the final refined model and map (dark-purple line) and a modified “scrambled” model (random displacement of all atoms by 0.5 Å), refined against 50% of the data, compared against half map 1 (solid purple line) or the independent half map 2 (dashed light-purple line). A thin dashed line indicates an FSC value of 0.5. **c** Two 90-degree related views of the helical reconstruction are shown from the side and the top, next to two slab views. **d** Two 180-degree related views of the two tetramers refined in isolation are visualized. The cryo-EM densities in (c,d) are colored according to local resolution, which was estimated using RELION-3.1 and visualized in UCSF ChimeraX (ref. ^3^). **e,f** The overall model to density fit between the final refined model (PDB-7P3R) and (e) the 4.1 Å cryo-EM map (EMD-13185-additional map 1) or (f) sections of the peripheral Head and Neck domains of the 3.7 Å map (EMD-13185). The individual copies of MakA are colored in shades of red and orange with the transmembrane domains highlighted blue. Selected bulky residues are indicated and labeled with amino acid code and residue number in red.

**Supplementary Table 1.**
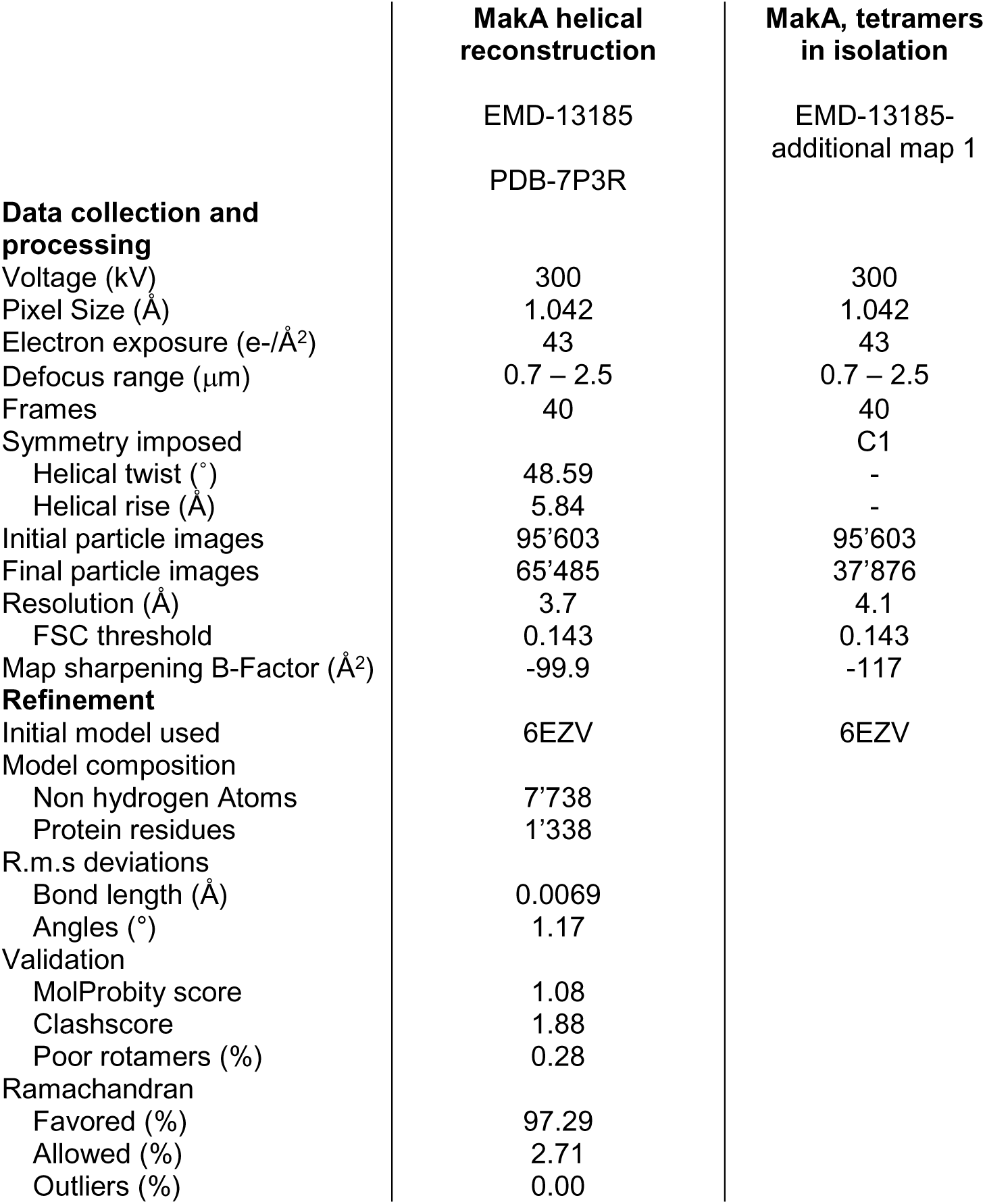
Cryo-EM data collection, refinement and model statistics.

